# Redundancy protects processing speed in healthy individuals with accelerated brain aging

**DOI:** 10.1101/2024.07.12.603314

**Authors:** William Stanford, Peter J. Mucha, Eran Dayan

## Abstract

Recent advancements in computational learning techniques have enabled the estimation of brain age (BA) from neuroimaging data. The difference between chronological age (CA) and BA, known as the BA gap, can potentially serve as a biomarker of brain health. Studies, however, have documented low correlations between BA gap and cognition in healthy aging. This suggests that protective mechanisms in the brain may help counter the effect of accelerated brain aging. Here, we investigated whether redundancy in brain networks may protect cognitive function in individuals with accelerated brain aging. First, we employed deep learning to estimate individual brain ages from structural magnetic resonance imaging (MRI). Next, we associated CA, BA, and BA gap, with cognitive measures and network topology derived from diffusion MRI and tractography. We found that CA and BA were both similarly related to cognitive measures and network topology, while BA gap did not show strong relationships in either domain. Despite observing no strong relationships between brain-age gap (BA gap) and demographic variables, cognitive measures, or topological features in healthy aging, individuals with accelerated aging (BA gap^+^) exhibited lower average degree and redundancy within the dorsal attention network compared to those with delayed aging (BA gap^-^). Furthermore, redundancy in the dorsal attention network was positively associated with processing speed in BA gap^+^ individuals. These results indicate a potential neuroprotective role of redundancy in structural brain networks for mitigating the impact of accelerated brain atrophy on cognitive performance in healthy aging.

## Introduction

Aging is a multi-facetted process associated with brain atrophy and decline in a wide range of cognitive functions.^1^ In recent years, advances in computational learning techniques have yielded data-driven methods for investigating differences in brain aging.^2^ In particular, predictions from models that estimate *brain age*^3,4^ (BA) from neuroimaging data enable researchers to study the disparity between an individual’s chronological age (CA) and biological age (BA), known as a *brain-age gap estimate*. Brain-age gap (BA gap) estimates have shown potential for clinical utility as a biomarker of brain health.^5,6^ For example, a positive BA gap (with BA > CA), indicating accelerated brain aging, has been linked to greater longitudinal physiological aging, age-related cognitive decline,^7,8^ conversion from mild cognitive impairment (MCI) to dementia,^9,10^ and mortality.^4,8^ While modifiable lifestyle factors have been shown to play a role in delaying and/or accelerating brain aging,^11–16^ genetics, and early life experiences, can also lead to accentuated brain aging.^17–21^ Interestingly, the magnitude of correlations reported between BA gap and cognition in healthy aging are typically low (r < 0.1),^5,22^ providing support for theories of cognitive aging.^23–28^ Thus brain-age studies in healthy aging populations provide an opportunity to study mechanisms that counter cognitive decline despite the presence of accelerated brain atrophy. As we develop a better understanding of the processes that lead to greater than expected brain aging, it’s critical to study mechanisms that could protect cognitive function in the presence of accelerated brain-atrophy in healthy aging.^22^

A mechanism that has shown promise as neuroprotective in cognitive aging is redundancy in brain networks.^29^ Redundancy protects systems from the failure of individual components; this principle is ubiquitous in biology,^29–35^ and in engineering.^36^ In the context of brain networks, redundancy can be defined as the number of direct and indirect paths between two brain regions.^37^ Duplicate pathways are believed to support robust information transmission between brain regions should some of the paths fail due to aging, disease processes, and/or environmental perturbations. In functional brain networks, redundancy has been shown to support executive function,^38^ and hippocampal redundancy has been found to be neuroprotective in healthy and pathological-aging.^39–41^ More recently, structural redundancy in cortical brain networks was discovered to mitigate age-related changes in the ability of brain networks to control their dynamics, and associate with processing speed in healthy aging.^42^ Incorporation of BA gaps into the study of redundancy in brain networks could enable us to understand how redundancy protects cognitive function before detectable differences in cognition occur.

In this study, we leverage deep learning to estimate BAs and BA gaps in healthy-aging participants of the HCP-Aging dataset.^43^ We trained the Simple Fully Convolutional Neural Network (SFCN) model^44^ to estimate the age of 451 participants via their anatomical MR images (Fig. 1A). This model was trained using the KL-divergence loss function, which enables a smooth decrease in the loss as prediction improves (Fig. 1B). Once model training was complete, we estimated the BAs of the remaining 193 participants in our dataset (Fig. 1C). We examine how demographics, topological measures of structural brain networks derived from diffusion MRI (dMRI), and several cognitive measures relate to BA and BA gaps (Fig. 1D). Then, we partition our participants into those with accelerated brain aging (BA gap^+^), and those with delayed brain aging (BA gap^-^) to investigate the role of redundancy in comparison with other major topological properties in supporting cognitive function in individuals with older than expected brain ages (Fig. 1E). We hypothesized that redundancy would serve as a neuroprotective mechanism in individuals with accelerated brain aging, mitigating the effect of accelerated brain-atrophy from impacting cognitive performance.

**Figure 1.**
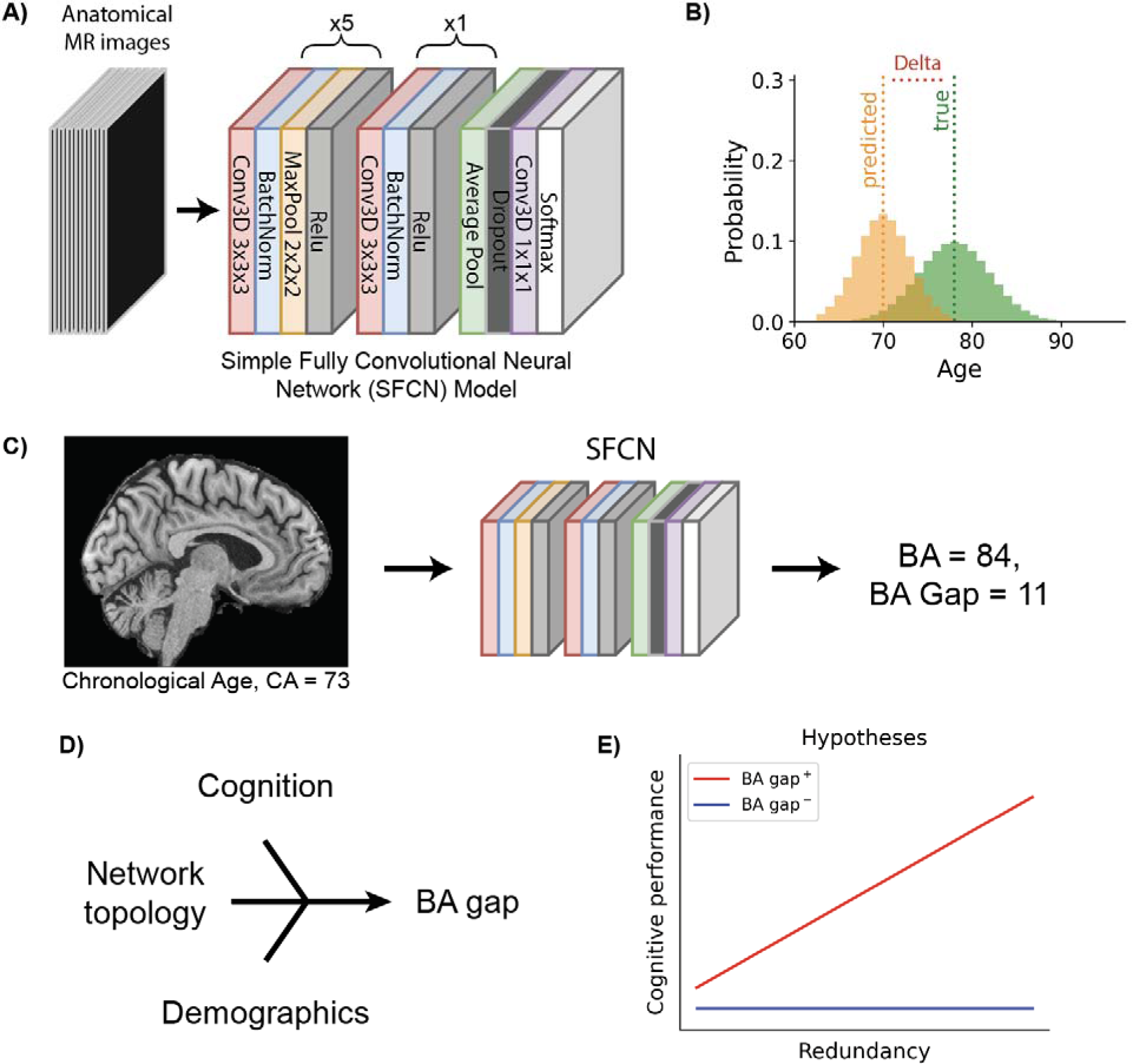
Study outline. **(A)** We first trained a neural network to predict an individual’s chronological age (CA) via 451 anatomical MR images using the Simple Fully Convolutional Neural Network (SFCN) model. **(B)** The model was trained via the KL-divergence loss function, using the probability distribution output by the model and a soft label in the form of a probability distribution centered on the ground-truth age. **(C)** Then we used the trained SFCN to estimate the brain ages (BAs) of the remaining 193 participants whose MR images were not used during training. An example of an individual with accelerated aging is illustrated above. Their chronological age (CA) was 73, and their BA was 84, giving a brain-age gap (BA gap) of +11. **(D)** Given estimates of BAs from all participants, we investigated the relationships between cognitive performance, topological measures of structural brain networks, demographics, and BA gap. **(E)** Next, we divided participants into those with accelerated brain aging (BA gap^+^), and delayed brain aging (BA gap^-^). We hypothesized that in individuals with accelerated brain aging, redundancy would be associated with greater levels of cognitive performance.

## Methods

### Participants

We used T1-weighted anatomical MR images and preprocessed dMRI data from participants in the Human Connectome Project – Aging database.^43^ Participants included were within the age range of 40-90, and classified as displaying typical aging according to the criteria outlined by the Human Connectome Project.^45^ All participants provided informed consent and all procedures had been pre-approved by local Institutional Review Boards.

### Deep learning model

Deep learning models have become increasingly used for estimating BA using neuroimaging data.^46^ In this study, we use the Simple Fully Convolutional Network (SFCN),^44^ a state-of-the-art BA prediction model that relies only on minimally preprocessed T1-weighted anatomical MR images. This model contains 7 blocks. The first 5 blocks each contain a 3×3×3 3D convolution layer, followed by a layer for batch normalization, then a 2×2×2 max pooling layer, and finally a ReLU activation layer. The 6^th^ block contains a 1×1×1 3D convolution layer, a batch normalization layer, and then ReLU activation. The 7^th^ block has an average pooling layer, followed by a layer for dropout (rate=0.5), a 1×1×1 3D convolution layer and, finally, a softmax. The final output layer contains 50 digits that are soft labels representing a predicted probability that the participant’s age is within a one-year period between the ages of 40-90. This final layer was modified to accommodate the larger age-range in our dataset. To calculate a predicted age for each participant, we use a weighted average of each age bin:

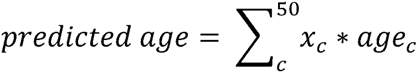

where *x_c_*is the model’s output for the probability of the *c*^th^ age bin, and *age_c_* is the center of the respective age interval. To train the model, we compute the Kullback-Leibler (KL) divergence loss using the soft labels (predicted probabilities) output by the final softmax layer.

### Model training and regularization

We used T1-weighted anatomical MRI images from 644 HCP-Aging participants with a 70/30 (451/193) train/test split stratified by the decades of participants’ ages. On the training data, we additionally performed stratified 5-fold cross-validation. We regularized the model via dropout (ratio=0.5), and performed data augmentation every epoch by randomly shifting the training data by 0, 1, or 2 voxels along each axis and mirroring images across the sagittal plane with 50% probability. We used a batch size 6 and trained for 300 epochs on an Nvidia Tesla V100 GPU with 16G memory. The model was implemented in PyTorch. The best performing model of the 5-fold cross-validation was selected to use on the held-out test set. For this model, we averaged the prediction of the 5 best epochs during training to a single BA prediction for each participant.

### Demographics, cognitive measures, and physical fitness

For demographic data, we used age and years of education. The primary cognitive measure we examined in our study was processing speed due to the replicated finding that processing speed is highly dependent on communication along white-matter tracts.^47,48^ To assess participants’ processing speed, they underwent the Pattern Comparison Processing Speed Test,^49^ where they were presented with two stimuli simultaneously and tasked to determine if the objects were the same. This task lasted 85 seconds, wherein participant scores were based on the number of correct judgements within this time window. We also utilized measures of episodic memory evaluated via the Rey Auditory and Verbal Learning Task (RAVLT),^50^ executive function determined by the Flanker Inhibitory Control and Attention Test^51^, and vocabulary comprehension tested with the Picture Vocabulary Test.^52^ Finally, we used MoCA scores as general measures of cognition.^53^ Previous studies have indicated that physical fitness is associated with brain maintenance.^15,16,54^ Therefore, we also examined relationships between BA gap and the physical fitness measure of walk endurance,^55^ where participants were asked to walk as far as they could within two minutes.

### Image acquisition and processing

We used T1-weighted anatomical MR images taken on a 3 Tesla Siemens Prisma Scanner. The sequence used was a multi-echo magnetization prepared rapid gradient echo (MPRAGE) sequence (voxel size: 0.8×0.8×0.8mm, TR = 2500ms, TE = 1.8/3.6/5.4/7.2ms, flip angle = 8 degrees). These images underwent a minimal preprocessing pipeline that included brain extraction, bias correction, and linear registration to the MNI152 space.^56^ We also made use of diffusion dMRI data acquired as part of the HCP-Aging project, with b-values of 1500s/mm^2^ and 3000s/mm^2^, 93 and 92 sampling directions, in-plane resolution of 1.5mm and a slice thickness of 1.5mm. We utilized preprocessed dMRI data according to the preprocessing pipeline provided by https://brain.labsolver.org/hcp_a.html. This pipeline included susceptibility artifact detection with TOPOP from the Tiny FSL package (http://github.com/frankyeh/TinyFSL), AC-PC alignment, restricted diffusion imaging,^57^ and generalized q-sampling.^58^ This preprocessing was done with resources at the Extreme Science and Engineering Discovery Environment (XSEDE)^59^ using the allocation TG-CIS200026.

### Network construction

To construct structural brain networks, we used preprocessed dMRI data reconstructed in DSI Studio (http://dsi-studio.labsolver.org). We used 5,000,000 streamlines while performing whole-brain fiber tracking, and registered these streamlines according to 400 brain regions in the Schaefer Local-Global cortical parcellation.^60^ This parcellation subdivides the human cortex into 17 large-scale networks based on functional subdivisions. Edge weights between two brain regions were defined as the number of streamlines passing through them. Edges were removed if the number of streamlines was below 0.001 of the maximum edge weight within each respective subject. Network metrics were computed with the binarized version of the structural networks, where edges with at least one streamline after thresholding were set to 1, and edges with no streamlines were set to 0.

### Topological measures

#### Redundancy

With our structural networks, we calculated redundancy as the number of direct and indirect non-circular paths between each pair of brain regions up to a specified length (*L* = 4).^37^ Redundancy between two nodes is defined as:

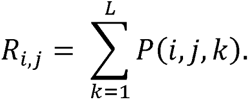

where *P*(*i, j, k*) is number of non-circular direct and indirect paths of length *k* between brain regions *i*, and *j,* as calculated with the *all_simple_paths* function in NetworkX.^61^ To compute redundancy for a single brain region *R_i_*, we summed across all node pairs *R_i,j_* for j from 1 to *n*, where *n* is the size of the parcellation. To compute redundancy within each of the 17 large-scale networks, we summed nodal redundancy for all brain regions within each respective network. For global redundancy, we summed the nodal redundancy of all brain regions in the parcellation. Because redundancy shows an exponential distribution across age,^38^ we performed all analyses with the logarithm of redundancy to linearize this feature.

#### Global and local efficiency

Measures of efficiency assume that communication on a network is inversely proportional to the shortest path between pairs of nodes. Global efficiency is the average inverse of the shortest path length between each pair of nodes,^62^ defined as:

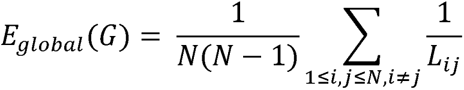

where *N* is the number of nodes in the graph *G*, and *L_ij_* is the length between nodes *i* and *j*. Local efficiency is calculated around a single node, as the average global efficiency on the subgraph induced by that node,^63^ defined below as:

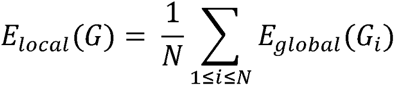

where *G_i_* is the set of all neighboring nodes of node *i* (but not including *i* itself). To compute local efficiency of each of the 17 large-scale networks, we averaged the local efficiencies of the nodes within each respective network. For average local efficiency of the entire network, we averaged the local efficiencies of all 400 nodes within the full brain network.

#### Degree

A node’s degree is defined as the number of nodes it is connected to. We computed the average degree in each participant’s structural brain network, as well as the average degrees of each of the 17 large-scale networks.

#### Max k-core

A *k*-core of a graph *G* is a connected subgraph of *G* where all nodes have at least degree *k* after iteratively deleting all nodes with degree less than *k.*^64^ We computed all the *k*-cores of each participant’s structural networks, and took the max *k*-core as a global topological metric, as well as the max *k*-core in each of the 17 large-scale networks.

### Statistical analysis

We performed group comparisons of several demographic, cognitive, and topological measures between individuals with older than expected, and younger than expected brains. We also calculated associations between these variables. To compare correlations across groups, we used the Fischer z-transformed correlations.^65^ We computed linear regressions between redundancy and cognitive measures, and to correct the bias in our BA model. Finally, we performed mediation analyses to investigate the extent to which network measures mediated the relationships between BA gap and cognition. When appropriate, we corrected for multiple comparisons accounting for the multiple cognitive scores, topological measures, and large-scale networks we assessed. Group comparisons were done with Welch’s T-tests. Associations were investigated via computing the Pearson’s correlation between the relevant variables. Welch’s T-tests, Pearson’s correlations, and mediation analysis were performed using the python package Pingouin.^66^ Linear regressions were computed in SciPy.^67^ To compute z-scored Pearson’s correlations we used the *corrstats.py* function at https://github.com/psinger/CorrelationStats/tree/master.

### Plotting

We created custom scripts for plotting and data visualization using the Matplotlib,^68^ Pandas,^69^ and Seaborn^70^ packages.

## Results

### Model performance and bias correction

We trained a modified version of the Simple Fully Convolutional Network (SFCN),^44^ a high-performing BA prediction model, on T1-weighted anatomical MR images from 70% of our HCP-Aging dataset for 300 epochs using stratified 5-fold cross-validation. This model was trained to predict a participant’s CA via features in their 3D T1-weighted anatomical MR image. After training, we used the best performing model to estimate the BAs of the remaining 30% of the dataset. For these estimations, we used an ensembling approach to improve model performance,^71^ in which we averaged the model’s predictions across the 5 best epochs during training. From these averaged predictions, we derived BA gap estimates, as the difference between BA and CA (BA gap = BA – CA). In association with CA, these BA gap estimates had a mean absolute error (MAE) of 4.79 ± 3.73, an *r* = 0.9 (Fig. 2A). Our model displayed an expected BA bias,^5^ in which BA gap showed a strong rank-correlation with CA (Spearman’s *ρ* = -0.59, *p* = 2.72e^-19^) (Fig. 2B). To alleviate this bias, we used a linear bias correction method commonly employed in many BA studies.^72^ After bias correction, our model achieved a MAE of 5.18 ± 4.21 years (Fig. 2C). The bias corrected estimated BAs, and BA gaps, were used in all following analyses.

**Figure 2.**
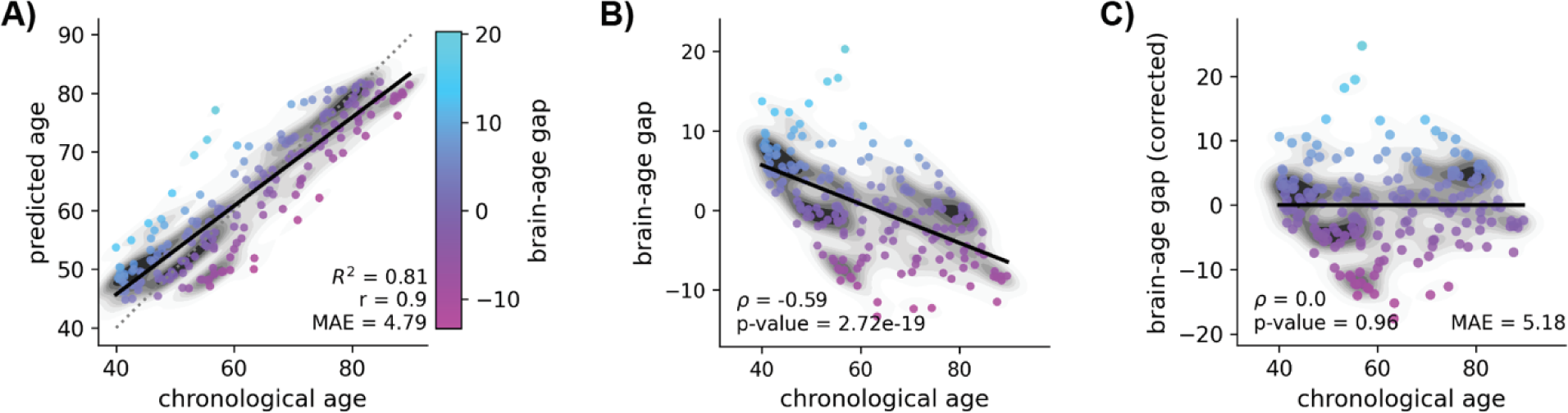
Model performance and bias correction. **(A)** Our model achieved a mean absolute error (MAE) of 4.79 ± 3.73 years on the age prediction task. **(B)** As shown in other brain-age models, our model’s performance of the SFCN was heavily biased by the imbalanced distribution of our training data. **(C)** After bias correction, our model’s performance dropped to an MAE of 5.18 ± 4.21 years.

### Brain age and chronological age are similarly correlated to cognitive scores and topological measures

After training our model, we evaluated the extent to which BA and CA differed in their associations with cognitive scores and topological measures of brain networks. For cognitive measures, we used measures of processing speed,^73^ episodic memory,^50^ executive function^74^ and vocabulary comprehension.^52^ We found that BA and CA showed highly similar associations with each of these cognitive domains (Fig. 3A and Table 1). In particular, episodic memory, processing speed, and executive function all showed expected negative relationships with BA and CA, while vocabulary comprehension showed an expected positive relationship with BA and CA. For topological measures, we associated measures of redundancy,^37^ along with local efficiency,^62,63^ average degree, and max k-core,^64^ with BA and CA. Each topological measure aside from local efficiency showed a significant negative relationship with both BA and CA (Fig. 3B and Table 2). While all of these correlations were moderately stronger for BA, none of the differences in correlation strengths were significant (Table 2). The difference between BA and CA (BA gap) was not significantly related with any of the cognitive scores evaluated (Fig. 3C), or topological measures assessed (Fig. 3D).

**Figure 3.**
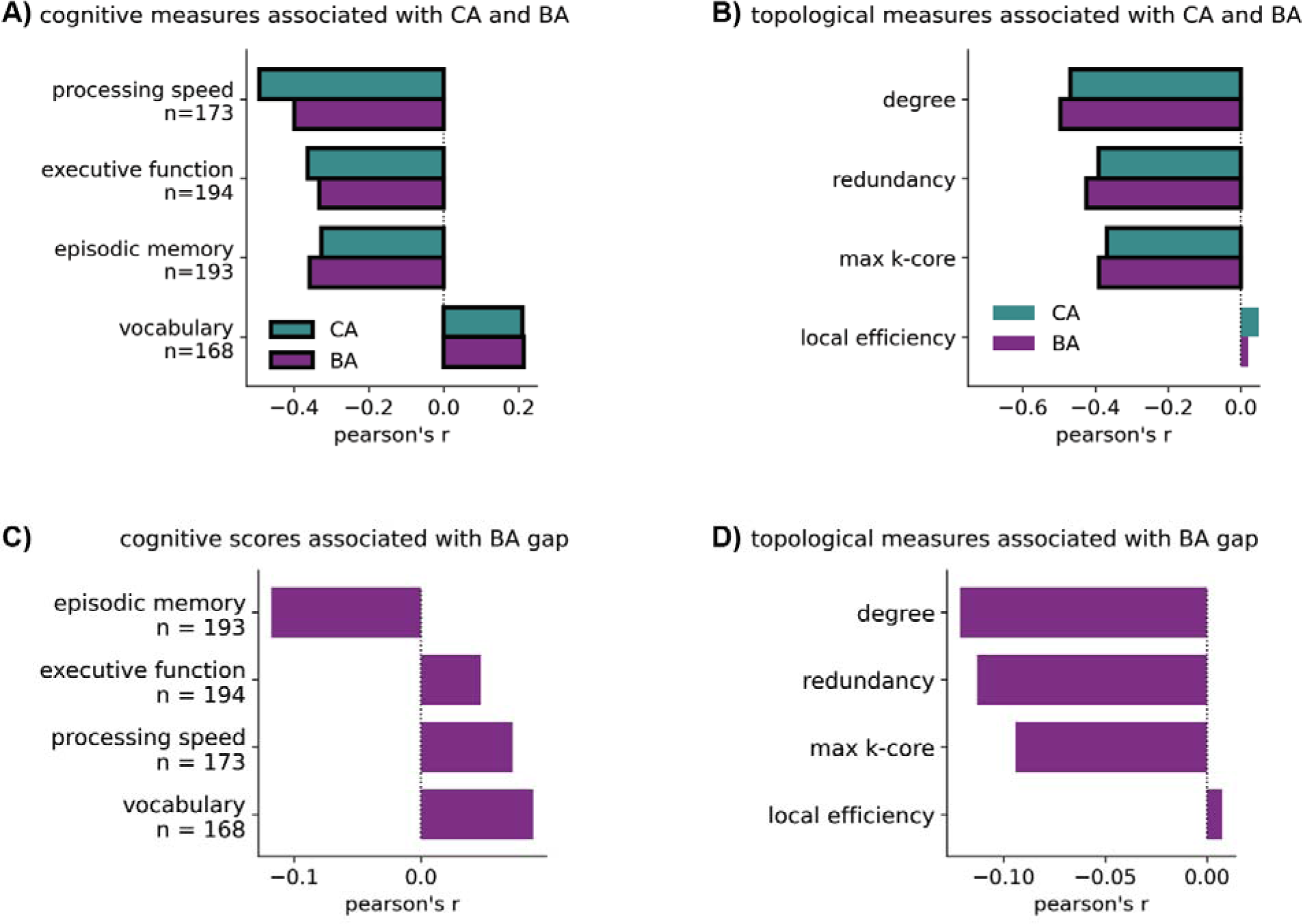
Chronological age (CA) and brain age (BA) show similar associations to cognitive scores and topological measures, but BA gap did not have significant relationships with these features in our sample. **(A)** CA and BA showed consistent relationships with the cognitive measures evaluated in our study. **(B)** Topological measures were also similarly related to both CA and BA. **(C)** The extent to which an individual’s estimated brain age deviated from their chronological age (BA gap) did not show any significant relationships with the cognitive measures used in our study. **(D)** For topological measures, we observed marginal negative relationships with BA gap. Significant correlations, if present, are indicated by black borders around the respective bars. In each panel we used the Bonferroni method to correct for multiple comparisons. For panels A, and C, we corrected for the four cognitive measures evaluated. In panels C, and D, we corrected for the 4 topological measures assessed. For each panel, the significance threshold was *p < 0.05/4.

**Table 1.** Cognitive scores show similar associations with CA and BA. Pearson’s *r*’s are reported for the relationships between each cognitive function, CA, and BA, respectively, as well as their z-scored differences.

| Topological measures associated with chronological age (CA) and brain age (BA) |  |  |  |  |  |  |
| --- | --- | --- | --- | --- | --- | --- |
| Cognitive scores | chronological age (CA) |  | brain-age (BA) |  | Differential correlations |  |
|  | r | p-value | r | p-value | z-scored difference | p-value |
| vocabulary<br>N=168 | 0.21 | 0.0062 | 0.21 | 5.40E-03 | 0.03 | 0.9737 |
| episodic memory<br>N=193 | -0.33 | 3.69E-06 | -0.36 | 3.36E-07 | 0.34 | 0.7301 |
| executive<br>function<br>N=194 | -0.36 | 1.93E-07 | -0.33 | 2.15E-06 | 0.34 | 0.7333 |
| processing<br>speed<br>N=173 | -0.49 | 6.29E-12 | -0.4 | 5.48E-08 | 1.07 | 0.2836 |

**Topological measures associated with chronological age (CA) and brain age (BA)**

**Table 2.** Topological measures show similar associations with CA and BA. Pearson’s *r*’s are reported for the relationships between each global topological measure, CA, and BA, respectively, as well as their z-scored differences.

| topological measures | chronological age (CA) |  | brain-age (BA) |  | Differential correlations |  |
| --- | --- | --- | --- | --- | --- | --- |
|  | r | p bonf. | r | p bonf. | z-scored difference | p bonf. |
| degree | -0.47 | 6.79E-12 | -0.50 | 2.37E-13 | 0.35 | 0.7268 |
| max k-core | -0.37 | 1.38E-07 | -0.39 | 2.01E-08 | 0.25 | 0.8012 |
| redundancy | -0.39 | 1.85E-08 | -0.42 | 7.90E-10 | 0.39 | 0.7002 |
| local efficiency | 0.05 | 0.50 | -0.54 | 7.91E-16 | 0.27 | 0.784 |

### Individuals with accelerated aging and delayed aging exhibit no differences in demographics, cognitive scores, or physical fitness

Next, we partitioned our subjects into BA gap^+^ and BA gap^-^ participants based on whether the magnitude of each participant’s BA deviated from their CA (Fig 4A). Individuals were considered BA gap^+^ if their BA was older than their CA by more than one year (BA – CA > 1, n=90). BA gap^-^ participants were those with younger than expected BAs at the same magnitude (BA – CA < -1, n=83). We found no differences between BA gap^+^ and BA gap^-^ participants in age (t_170.9_ = -0.47, *p_bonf_*. = 0.995) (Fig. 4B.i), years of education (t_168.8_ = 0.09, *p_bonf._* = 0.995) (Fig. 4B.ii), or MoCA scores (t_166.1_ = -0.29, *p_bonf._* = 0.995) (Fig. 4B.iii). We also found no differences between these two groups for processing speed (t_151.0_ = 0.32, *p_bonf._* = 0.995), episodic memory (t_169.0_ = -1.41, *p_bonf._* = 0.572), executive function (t_162.1_ = -0.32, *p_bonf._* = 0.995), or vocabulary comprehension (t_137.9_ = 1.20, *p_bonf_*. = 0.920) (Fig. 4C). We then compared the BA gap^+^ and BA gap^-^ groups using other threshold magnitudes for group definition of *k* = [0, 1, 2, 3, 4, 5 years] where participants were BA gap ^+^ if BA – CA > *k*, and BA gap ^-^ if BA – CA < -*k*. We did not observe any group differences for these demographic variables, or cognitive measures when considering these alternate thresholds used to partition into BA gap^+^ and BA gap^-^ participants (Tables S1-S7). We also did not observe any differences in levels of physical fitness between BA gap^+^ and BA gap^-^ groups (Table S8).

**Figure 4.**
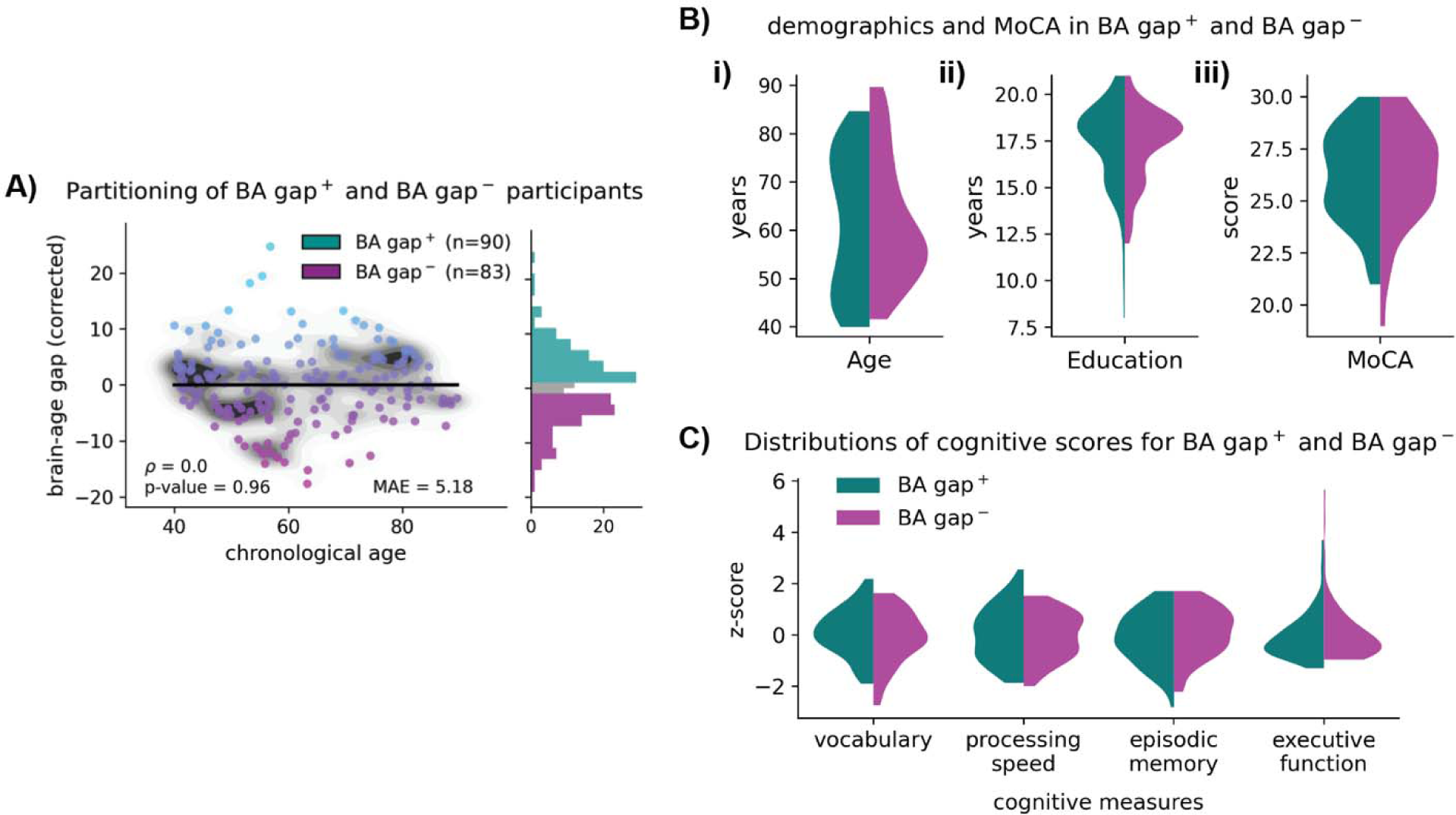
Individuals with accelerated aging (BA gap^+^) and delayed aging (BA gap^-^) show no differences in demographic variables or cognitive measures. **(A)** Participants were divided into two groups based on the extent to which their brain age deviated from their chronological age. BA gap**^+^** were participants with accelerated aging (BA gap > 1), while BA gap**^-^** were participants with delayed aging (BA gap < -1). **(B)** BA gap**^+^** and BA gap**^-^** participants showed no group level differences in chronological age (i), years of education (ii), or MoCA scores (iii). **(C)** z-scored cognitive scores were also consistent between BA gap**^+^** and BA gap**^-^** participants. Welch’s T-tests were used in each comparison. In panel C we used the Bonferroni method to correct for multiple comparisons for the 4 cognitive measures evaluated, which set the significance threshold to *p < 0.05/4.

### Individuals with accelerated brain aging exhibit differences in global topological measures of brain networks

Following our investigation into demographic and cognitive differences between BA gap^+^ and BA gap^-^ participants, we compared the aforementioned global topological measures of brain networks between these two groups (Fig. 5A). The structural brain networks showed similar average local efficiency in BA gap^+^ and BA gap^-^ participants (t_164.4_ = -0.59, *p_bonf._* = 0.552) and lower max degree in BA gap^+^ (t_165.5_ = -2.69, *p_bonf._* = 0.0314) (Fig. 5A.ii). We also observed marginal differences in max k-core (t_161.8_ = -2.27, *p_bonf._* = 0.0979) (Fig. 5A.iii), and total network redundancy (t_165.5_ = -2.52, *p_bonf._*= 0.0514) (Fig. 5A.iv), but these differences were not significant after correcting for multiple comparisons. The lack of difference in average local efficiency between BA gap^+^ and BA gap^-^ participants was consistent across thresholds assessed (Table S9). For degree, we only observed significant differences with *k* = [1, 4] (Table S10). Max k-core did not show any significant differences across the thresholds examined (Table S11). Total redundancy trended towards significance for *k* = [1, 4] (Table S12).

**Figure 5.**
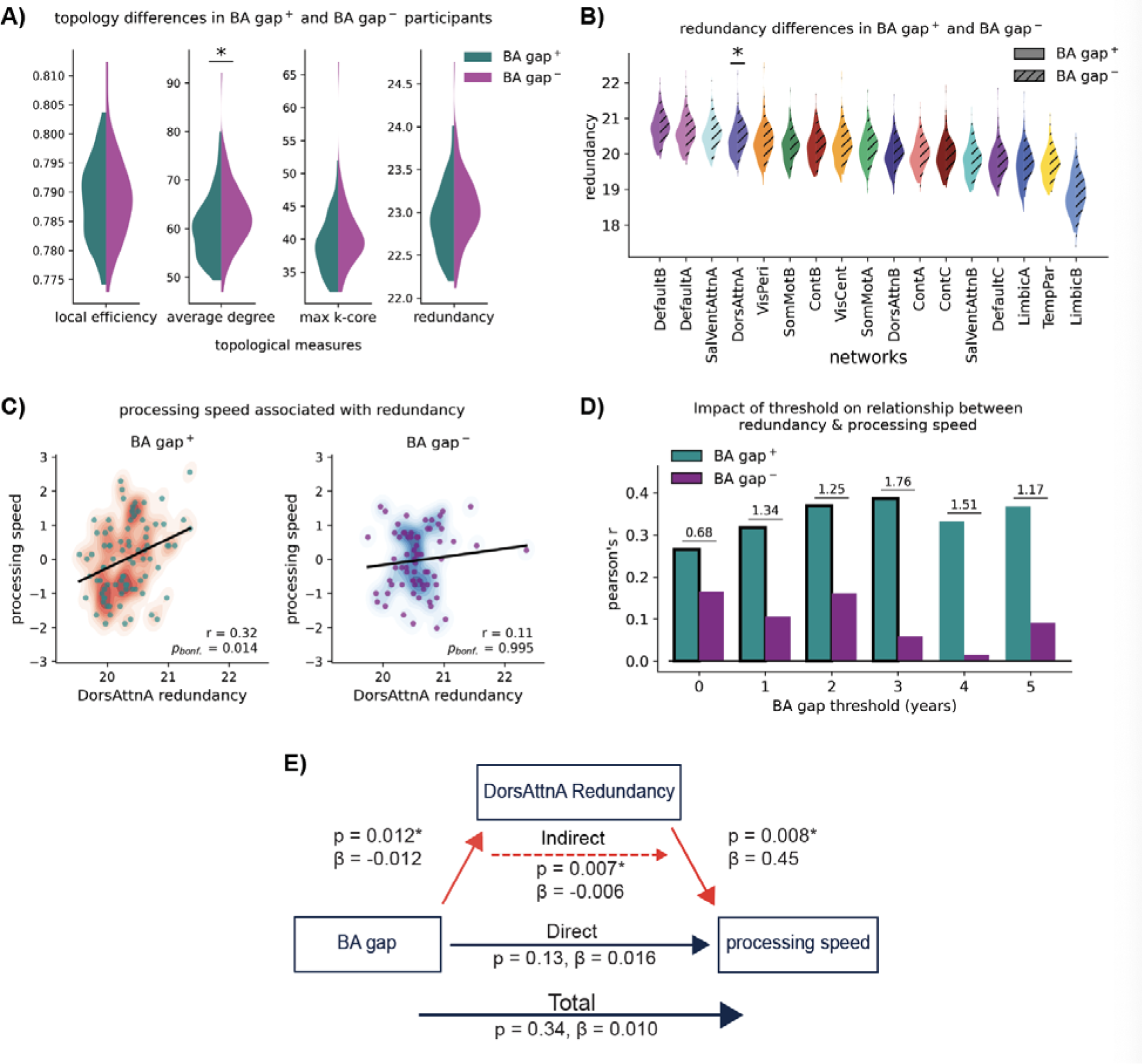
Redundancy supports processing speed in individuals with accelerated aging. **(A)** The brain networks of BA gap^+^ participants had lower average degree than BA gap^-^ participants. These groups did not differ in their average local efficiency, max *k*-cores, or total redundancy. **(B)** BA gap^+^ participants exhibited lower levels of redundancy in the dorsal attention network (DorsAttnA). **(C)** Processing speed was positively associated with redundancy in the dorsal attention network (DorsAttnA) for BA gap^+^ participants, but not for BA gap^-^ participants. **(D)** The relationship in panel C became stronger when considering individuals with increasingly accelerated aging. Significant correlations, if present, are indicated by black borders around the respective bars. The z-scored differences between the correlations observed in BA gap^+^ and BA gap^-^ participants are displayed above each pair of bars at the respective threshold levels**. (E)** Redundancy in the dorsal attention network (DorsAttnA) positively mediated the relationship between BA gap and processing speed. The Bonferroni method to correct for multiple comparisons was used to detect significant relationships in each panel. In panel A we corrected for comparisons across 4 global topological measures (*p < 0.05/4). In panel B we corrected for comparisons across 17 networks within our parcellation (*p < 0.05/17). In panel’s C, D, and E, we corrected for comparisons across 4 cognitive measures (*p < 0.05/4), *p_bonf._* indicates already corrected p-values where *p_bonf._* = *p*\*(*number of comparisons*).

### Individuals with accelerated aging have less redundancy in the dorsal attention network

Motivated by previous studies indicating the neuroprotective effect of redundancy in brain networks,^38–42^ we further examined group level differences in redundancy in each of the 17 large-scale networks in the local-global Schaefer-Yeo parcellation.^60^ BA gap^+^ participants had less redundancy in each network (Fig. 5B and Table 3). However, these differences were only significant within the dorsal attention network after correction for multiple comparisons (DorsAttnA, t_165.76_ = -3.33, *p_bonf._* = 0.0180). This relationship persisted when *k >* 1 was used to partition BA gap^+^ and BA gap^-^ participants (Table S13).

**Table 3.** BA gap^+^ participants had less redundancy in the dorsal attention network (DorsAttnA). Welch’s T-tests were used in each comparison. We corrected for multiple comparisons across 17 large-scale networks using the Bonferroni method, *p_bonf._* indicates already corrected p-values where *p_bonf._* = *p*\*17.

| Redundancy differences in large-scale networks |  |  |  |  |  |
| --- | --- | --- | --- | --- | --- |
| | BA gap+<br>(mean ± std) | BA gap-<br>(mean ± std) | T | dof | p bonf. |
| DefaultB | 20.68 ± 0.39 | 20.79 ± 0.40 | -1.87 | 168.37 | 0.995 |
| DefaultA | 20.54 ± 0.40 | 20.67 ± 0.43 | -2.01 | 165.64 | 0.777 |
| SalVentAttnA | 20.46 ± 0.38 | 20.61 ± 0.40 | -2.43 | 166.54 | 0.273 |
| **DorsAttnA\*** | 20.33 ± 0.40 | 20.54 ± 0.43 | -3.33 | 165.76 | 0.018 |
| VisPeri | 20.22 ± 0.43 | 20.41 ± 0.47 | -2.65 | 163.98 | 0.149 |
| SomMotB | 20.14 ± 0.40 | 20.28 ± 0.39 | -2.34 | 168.97 | 0.347 |
| ContB | 20.14 ± 0.42 | 20.29 ± 0.41 | -2.47 | 169.19 | 0.248 |
| VisCent | 20.12 ± 0.43 | 20.30 ± 0.45 | -2.75 | 167.20 | 0.113 |
| SomMotA | 20.09 ± 0.45 | 20.25 ± 0.43 | -2.39 | 169.65 | 0.304 |
| DorsAttnB | 19.92 ± 0.42 | 20.10 ± 0.41 | -2.88 | 168.98 | 0.077 |
| ContA | 19.90 ± 0.40 | 20.06 ± 0.41 | -2.49 | 166.92 | 0.234 |
| ContC | 19.89 ± 0.42 | 20.03 ± 0.43 | -2.12 | 167.16 | 0.608 |
| SalVentAttnB | 19.73 ± 0.43 | 19.87 ± 0.42 | -2.05 | 169.40 | 0.715 |
| DefaultC | 19.68 ± 0.40 | 19.84 ± 0.45 | -2.49 | 162.75 | 0.232 |
| LimbicA | 19.59 ± 0.50 | 19.79 ± 0.47 | -2.68 | 169.84 | 0.137 |
| TempPar | 19.60 ± 0.41 | 19.76 ± 0.42 | -2.50 | 167.36 | 0.230 |
| LimbicB | 18.81 ± 0.52 | 18.92 ± 0.55 | -1.37 | 166.36 | 0.995 |

### Redundancy in the dorsal attention network promotes processing speed in individuals with accelerated aging

After we identified lower levels of dorsal attention network redundancy in individuals with older than expected BA, we investigated the extent to which redundancy was associated with cognitive function. We found that redundancy in the dorsal attention network was positively associated with processing speed in BA gap^+^ participants (DorsAttnA, r = 0.32, *p_bonf._* = 0.014), but not in BA gap^-^ participants (DorsAttnA, r = 0.11, *p_bonf._* = 0.995) (Fig. 5C). Similar relationships were observed when using BA gap thresholds of 0, 1, 2, & 3 years (Fig. 5D and Table 4). Furthermore, the z-scored differences in correlations between BA gap^+^ and BA gap^-^ participants appeared to increase with threshold magnitude, suggesting an increased role for redundancy as the magnitude of the accelerated aging increased. For other cognitive functions, we saw positive trends between dorsal attention (DorsAttnA) redundancy and episodic memory for both BA gap^+^ and BA gap^-^ participants, but fewer of these relationships were significant across thresholds (Table S14). Neither executive function nor vocabulary showed significant relationships with dorsal attention redundancy (Table S15-S16). Finally, we performed mediation analyses to examine the influence of redundancy on the relationships between BA gap and the cognitive measures studied here. We found that dorsal attention redundancy had a significant indirect effect on the relationship between BA gap and processing speed (DorsAttnA, β = -0.006, *p* = 0.007, *p_bonf._* = 0.035) (Fig. 5E). We also observed a significant indirect effect of redundancy on the relationship between BA gap and episodic memory (β = -0.019, *p* = 0.0094, *p_bonf._* = 0.0376) (Table S17). However, dorsal attention redundancy did not mediate these relationships for executive function (DorsAttnA, β = 0.0279, p-value = 0.7114, *p_bonf_*. = 0.995) (Table S18), or vocabulary (DorsAttnA, β = 0.0471, *p* = 0.0336, *p_bonf._* = 0.1344) (Table S19).

**Table 4.** BA gap^+^ participants showed stronger relationships between dorsal attention network (DorsAttnA) redundancy and processing speed as deviation from expected age increased. We corrected for multiple comparisons across 4 cognitive measures assessed using the Bonferroni method, *p_bonf._*indicates already corrected p-values where *p_bonf._* = *p*\*4.

| BA gap<br>threshold<br>(years) | BA gap+ |  |  | BA gap- |  |  | Differential correlations |  |
| --- | --- | --- | --- | --- | --- | --- | --- | --- |
|  | n | r | p bonf. | n | r | p bonf. | z-scored<br>difference | p bonf. |
| 0 | 95 | 0.265* | 0.0382 | 78 | 0.165 | 0.6006 | 0.68 | 0.498 |
| 1 | 83 | 0.317* | 0.0139 | 70 | 0.106 | 0.995 | 1.34 | 0.179 |
| 2 | 68 | 0.368* | 0.0080 | 63 | 0.161 | 0.8296 | 1.25 | 0.211 |
| 3 | 57 | 0.386* | 0.0121 | 51 | 0.058 | 0.995 | 1.76 | 0.079 |
| 4 | 46 | 0.332 | 0.0969 | 43 | 0.014 | 0.995 | 1.51 | 0.132 |
| 5 | 39 | 0.367 | 0.0858 | 31 | 0.090 | 0.995 | 1.17 | 0.242 |

### Local efficiency in the dorsal attention network does not promote processing speed in individuals with accelerated aging

Our results thus far indicate that redundancy could support processing speed in individuals with accelerated aging. Redundancy counts the total number of direct and indirect paths between brain regions. We hypothesized that redundancy would support processing speed as it provides additional robustness in the communicative abilities between brain regions should connections fail. It’s also possible that efficient communication could be important for this cognitive function. Therefore, in following previous research,^41^ we computed the local efficiency of each brain region, and investigated the importance of average local efficiency in the dorsal attention network for processing speed. First, we compared the average local efficiency of each of the 17 large-scale networks in BA gap^+^ and BA gap^-^ participants (Fig. S1A and Table S20). We did not see significant differences in local efficiency for any of these networks between groups. Next, we associated processing speed with local efficiency of the dorsal attention network in BA gap^+^ and BA gap^-^ participants using several different thresholds for group definition (Fig. S1B). We did not find any significant relationships in either group regardless of the thresholds investigated. Finally, we investigated if local efficiency of the dorsal attention network mediated the relationship between BA gap and processing speed (Fig. S1C). While we did observe a significant relationship between BA gap and local efficiency (β = 0.0005, *p* = 0.0001), the indirect effect of local efficiency on the relationship between BA gap and processing speed was not significant (β = -0.003, *p* = 0.283).

## Discussion

We used deep learning techniques to estimate an individual’s BA via T1-weighted anatomical MR images. We found that BA and CA showed similar correlations for both cognition and network topology. Next, we observed that differences between an individual’s BA and CA (BA gap) in healthy aging did not show strong relationships with any of the demographic variables, cognitive measures, or topological features we examined. We then partitioned participants into those that exhibited accelerated aging (BA gap^+^), and individuals with delayed aging (BA gap^-^). These groups exhibited no differences in demographic variables or cognitive scores. However, the structural brain networks of BA gap^+^ individuals showed less overall average degree, as well as decreased redundancy within the dorsal attention network.

Furthermore, in individuals with accelerated aging, redundancy in the dorsal attention network was positively associated with processing speed, a relationship not present in those with delayed aging. These relationships were specific to processing speed, and not observed with other cognitive functions. Finally, redundancy in the dorsal attention network, but not local efficiency, mediated the relationships between BA gap and processing speed.

### Brain-age gap, demographic variables, and cognition

Education and physical fitness have both previously been linked to a younger than expected BA,^13,16,54^ supporting their hypothesized roles in cognitive reserve.^26,75–77^ These results would lead us to expect that BA gap^+^ individuals would show lower levels of physical fitness, and less educational attainment than BA gap^-^ participants. However, our results did not indicate significant differences in these demographics variables between BA gap^+^ and BA gap^-^ participants. For education, our results could be due to a lack of robustness of education as a measure of reserve.^22,78^ In the context of physical fitness, BA gap^-^ individuals did generally exhibit higher levels of fitness than those with BA gap^+^ participants, following the expected trend, but this effect was not significant in our sample. Future studies should utilize larger sample sizes and/or more sensitive measures to clarify the impact of education and physical fitness on BA gap. The cognitive measures we evaluated showed moderate relationships with BA gap, and no significant differences between BA gap^+^ and BA gap^-^ participants. We expect that this is primarily due to our focus on typically-aging participants, in which reported relationships between BA gap and cognition are low.^5,22^ However, several studies have reported stronger correlations than what we observed.^7,8,79^ Therefore, more work should be done in this area to clarify what relationships exist between BA gap and cognitive performance in typical aging.

### Age, brain-age gap, and topological metrics

We found that each of the global topological metrics we considered showed a negative relationship with age, which follows other reports of age-associated breakdowns in the efficiency^80,81^ and connectedness^82–84^ of structural brain networks. While we did not observe significant associations between BA gap and any topological measure, we did observe group level differences between BA gap^+^ and BA gap^-^ individuals. BA gap^+^ participants showed less redundancy and average degree, which, when taken together, suggests that a greater than expected BA is linked to an acceleration of age-related network atrophy.

### Neuroprotective effect of dorsal attention network redundancy in processing speed and episodic memory

We observed no differences in the cognitive function of BA gap^+^ and BA gap^-^ participants. We hypothesized that redundancy would partially protect cognitive function in individuals with accelerated brain-atrophy. For processing speed, our results indicate that redundancy in the dorsal attention network (DorsAttnA) may provide a neuroprotective effect that minimizes an older than expected BA from impacting cognitive function in healthy aging. For those with younger than expected brains, we observed a plateauing effect, where further increases in redundancy were not correlated with greater processing speed, similar to the results reported for hippocampal redundancy.^41^ The dorsal attention network is critical for visual processing speed,^85^ and age-associated disruptions to this network have been linked to age-related declines in processing speed.^42,86,87^ In addition, the dorsal attention network’s hub-like properties appear to be key to supporting the flow of information to the network during task-based activities.^88^ In this context, the neuroprotective effect we observed could be due to the additional robustness that redundancy provides for information flow to and from this network. We note that our results could not be explained by local efficiency of the dorsal attention network, implying that robust information flow supported via redundancy is more important for processing speed than efficient information flow along the shortest path. Our primary focus in this study was processing speed, as this function has shown strong relationships with measures of white matter connectivity.^47,48^ However, we also observed a significant mediation by redundancy between the BA gap and episodic memory relationship,^40^ suggesting that structural redundancy could play a role in this cognitive function too.

### Network topology and brain reserve

Topological measures of brain networks have frequently been studied as mechanisms that could support cognitive function in aging^89^ and neurodegenerative disease.^90–92^ In particular, redundancy has been studied in the context of healthy aging^38,40,41^ as well as in individuals with MCI.^40,41^ As numerous studies have indicated, topological properties appear to have the strongest impact on cognitive reserve in the early stages of advanced aging^93^ or AD.^40,90,91^ However, our study indicates that topological properties could protect cognitive function from accelerated biological aging before detectable differences in cognition are present, and that these properties can be isolated with the combination of deep learning and graph-based topological measures of brain networks.

### Future directions and limitations

Brain age in middle age has been related to accelerated aging and longitudinal cognitive decline.^7^ Future studies should evaluate the extent to which redundancy moderates these effects on cognitive function. Furthermore, we encourage future studies that mix BA prediction with topological analysis of brain networks, as these studies could provide key insight into the topological properties most relevant for cognitive reserve in aging and neurodegenerative diseases. The deep learning model we used was unimodal. A multimodal model could have performed better at our task, yielding a more accurate estimation of BA. However, models trained with only T1-weighted anatomical MR images appear to exhibit similar effect sizes to those that multimodal models capture in healthy aging.^5^ Another important consideration is the predictive accuracy of our BA model. We used the SFCN,^44^ with which we achieved an MAE comparable to current state-of-the-art results for similar dataset sizes and participant age-ranges.^46^ However, it is possible that using a different architecture, or dataset, such as UK Biobank,^6^ would have yielded a more accurate MAE. The impact these choices had on our results is difficult to estimate, but the magnitude of the observed associations between BA gap and cognitive function fell within the expected range when compared to similar studies.^5^ Finally, our primary focus in this study was processing speed, as processing speed has shown strong relationships with measures of white matter connectivity.^47,48^ However, we also observed a significant mediation by redundancy in the relationship between BA gap and episodic memory, suggesting that redundancy could play a neuroprotective role in this cognitive function too.^40^ Future studies should explore other topological metrics and methods for constructing brain networks in attempt to explain the mechanisms that enable individuals with accelerated brain aging to maintain levels of cognitive function similar to their peers.

### Conclusion

In conclusion, this study has provided a method to investigate how network mechanisms mitigate accelerated brain aging from impacting cognitive function in healthy aging. Our results indicate that a greater than expected BA may reflect age-related network degeneration, but is not always linked to detectable differences in cognition. Importantly, we discovered that individuals with accelerated brain aging could leverage structural redundancy in the dorsal attention network to preserve processing speed abilities despite experiencing greater brain atrophy than their peers. These data provide further evidence of the neuroprotective role of redundancy within brain networks to support cognitive function in aging.

## Conflicts of interest

None disclosed.

## Supporting information

Stanford_SM

## Acknowledgments

The study was supported by the National Institute On Aging of the National Institutes of Health under Award Number R01AG062590. The content is solely the responsibility of the authors and does not necessarily represent the official views of the National Institutes of Health.

## Notes

### Competing Interest Statement

The authors have declared no competing interest.

## Reference List

1 Novotný, J. S. et al. Physiological pattern of cognitive aging. medRxiv (2021).

2 Singh, N. M. et al. How machine learning is powering neuroimaging to improve brain health. Neuroinformatics 20, 943–964 (2022).

3 Franke, K. & Gaser, C. Ten years of BrainAGE as a neuroimaging biomarker of brain aging: what insights have we gained? Frontiers in neurology, 789 (2019).

4 Cole, J. H. & Franke, K. Predicting age using neuroimaging: innovative brain ageing biomarkers. Trends in neurosciences 40, 681–690 (2017).

5 Jirsaraie, R. J. et al. A systematic review of multimodal brain age studies: Uncovering a divergence between model accuracy and utility. Patterns 4 (2023).

6 Cole, J. H. Multimodality neuroimaging brain-age in UK biobank: relationship to biomedical, lifestyle, and cognitive factors. Neurobiology of aging 92, 34–42 (2020).

7 Elliott, M. L. et al. Brain-age in midlife is associated with accelerated biological aging and cognitive decline in a longitudinal birth cohort. Molecular psychiatry 26, 3829–3838 (2021).

8 Cole, J. H. et al. Brain age predicts mortality. Molecular psychiatry 23, 1385–1392 (2018).

9 Franke, K. & Gaser, C. Longitudinal changes in individual BrainAGE in healthy aging, mild cognitive impairment, and Alzheimer’s disease. GeroPsych (2012).

10 Gaser, C. et al. BrainAGE in mild cognitive impaired patients: predicting the conversion to Alzheimer’s disease. PloS one 8, e67346 (2013).

11 Luders, E., Cherbuin, N. & Gaser, C. Estimating brain age using high-resolution pattern recognition: Younger brains in long-term meditation practitioners. Neuroimage 134, 508–513 (2016).

12 Linli, Z., Feng, J., Zhao, W. & Guo, S. Associations between smoking and accelerated brain ageing. Progress in Neuro-Psychopharmacology and Biological Psychiatry 113, 110471 (2022).

13 Steffener, J. et al. Differences between chronological and brain age are related to education and self-reported physical activity. Neurobiology of aging 40, 138–144 (2016).

14 Kaplan, A. et al. The effect of a high-polyphenol Mediterranean diet (Green-MED) combined with physical activity on age-related brain atrophy: the Dietary Intervention Randomized Controlled Trial Polyphenols Unprocessed Study (DIRECT PLUS). The American Journal of Clinical Nutrition 115, 1270–1281 (2022).

15 Boraxbekk, C.-J., Salami, A., Wåhlin, A. & Nyberg, L. Physical activity over a decade modifies age-related decline in perfusion, gray matter volume, and functional connectivity of the posterior default-mode network—A multimodal approach. Neuroimage 131, 133–141 (2016).

16 Voss, M. W. et al. Fitness, but not physical activity, is related to functional integrity of brain networks associated with aging. Neuroimage 131, 113–125 (2016).

17 Kaufmann, T. et al. Genetics of brain age suggest an overlap with common brain disorders. BioRxiv, 303164 (2018).

18 Drobinin, V. et al. The developmental brain age is associated with adversity, depression, and functional outcomes among adolescents. Biological Psychiatry: Cognitive Neuroscience and Neuroimaging 7, 406–414 (2022).

19 Leonardsen, E. H. et al. Genetic architecture of brain age and its casual relations with brain and mental disorders. medRxiv, 2023.2001. 2009.23284310 (2023).

20 Wen, J. et al. The Genetic Heterogeneity of Multimodal Human Brain Age. bioRxiv, 2023.2004. 2013.536818 (2023).

21 Hedderich, D. M. et al. Increased brain age gap estimate (BrainAGE) in young adults after premature birth. Frontiers in Aging Neuroscience, 158 (2021).

22 Anatürk, M. et al. Prediction of brain age and cognitive age: Quantifying brain and cognitive maintenance in aging. Human brain mapping 42, 1626–1640 (2021).

23 Stern, Y. Cognitive reserve in ageing and Alzheimer’s disease. The Lancet Neurology 11, 1006–1012 (2012).

24 M Tucker, A. & Stern, Y. Cognitive reserve in aging. Current Alzheimer Research 8, 354–360 (2011).

25 Stern, Y. et al. Whitepaper: Defining and investigating cognitive reserve, brain reserve, and brain maintenance. Alzheimer’s & Dementia 16, 1305–1311 (2020).

26 Scarmeas, N. & Stern, Y. Cognitive reserve and lifestyle. Journal of clinical and experimental neuropsychology 25, 625–633 (2003).

27 Stern, Y. Cognitive reserve. Neuropsychologia 47, 2015–2028 (2009).

28 Stern, Y., Barnes, C. A., Grady, C., Jones, R. N. & Raz, N. Brain reserve, cognitive reserve, compensation, and maintenance: operationalization, validity, and mechanisms of cognitive resilience. Neurobiology of aging 83, 124–129 (2019).

29 Glassman, R. B. An hypothesis about redundancy and reliability in the brains of higher species: Analogies with genes, internal organs, and engineering systems. Neuroscience & Biobehavioral Reviews 11, 275–285 (1987).

30 Navlakha, S., He, X., Faloutsos, C. & Bar-Joseph, Z. Topological properties of robust biological and computational networks. Journal of the Royal Society Interface 11, 20140283 (2014).

31 Kafri, R., Springer, M. & Pilpel, Y. Genetic redundancy: new tricks for old genes. Cell 136, 389–392 (2009).

32 Pitkow, X. & Angelaki, D. E. Inference in the brain: statistics flowing in redundant population codes. Neuron 94, 943–953 (2017).

33 Lawton, J. & Brown, V. (Berlin, Springer, 1993).

34 Nyström, M. Redundancy and response diversity of functional groups: implications for the resilience of coral reefs. AMBIO: A Journal of the Human Environment 35, 30–35 (2006).

35 Cárdenas, A. et al. Greater functional diversity and redundancy of coral endolithic microbiomes align with lower coral bleaching susceptibility. The ISME journal 16, 2406–2420 (2022).

36 Billinton, R. & Allan, R. N. Reliability evaluation of engineering systems. (Springer, 1992).

37 Di Lanzo, C., Marzetti, L., Zappasodi, F., De Vico Fallani, F. & Pizzella, V. Redundancy as a graph-based index of frequency specific MEG functional connectivity. Computational and mathematical methods in medicine 2012 (2012).

38 Sadiq, M. U., Langella, S., Giovanello, K. S., Mucha, P. J. & Dayan, E. Accrual of functional redundancy along the lifespan and its effects on cognition. Neuroimage 229, 117737 (2021).

39. Langella, S. Functional Hippocampal Redundancy as a Measure of Resilience to Pathological Aging, The University of North Carolina at Chapel Hill, (2021).

40 Langella, S., Mucha, P. J., Giovanello, K. S., Dayan, E. & Initiative, A. s. D. N. The association between hippocampal volume and memory in pathological aging is mediated by functional redundancy. Neurobiology of Aging 108, 179–188 (2021).

41 Langella, S., Sadiq, M. U., Mucha, P. J., Giovanello, K. S. & Dayan, E. Lower functional hippocampal redundancy in mild cognitive impairment. Translational psychiatry 11, 1–12 (2021).

42 Stanford, W., Mucha, P. J. & Dayan, E. Age-related differences in network controllability are mitigated by redundancy in large-scale brain networks. Communications Biology 7, 701 (2024).

43 Harms, M. P. et al. Extending the Human Connectome Project across ages: Imaging protocols for the Lifespan Development and Aging projects. Neuroimage 183, 972–984 (2018).

44 Peng, H., Gong, W., Beckmann, C. F., Vedaldi, A. & Smith, S. M. Accurate brain age prediction with lightweight deep neural networks. Medical image analysis 68, 101871 (2021).

45 Bookheimer, S. Y. et al. The lifespan human connectome project in aging: an overview. Neuroimage 185, 335–348 (2019).

46 Tanveer, M. et al. Deep learning for brain age estimation: A systematic review. Information Fusion (2023).

47 Bullmore, E. & Sporns, O. The economy of brain network organization. Nature reviews neuroscience 13, 336–349 (2012).

48 Lynn, C. W. & Bassett, D. S. The physics of brain network structure, function and control. Nature Reviews Physics 1, 318–332 (2019).

49 Carlozzi, N. E. et al. NIH toolbox cognitive battery (NIHTB-CB): the NIHTB pattern comparison processing speed test. Journal of the International Neuropsychological Society 20, 630–641 (2014).

50 Rey, A. L’examen psychologique dans les cas d’encéphalopathie traumatique.(Les problems.). Archives de psychologie (1941).

51 Zelazo, P. D., et al. II. NIH Toolbox Cognition Battery (CB): Measuring executive function and attention. Monographs of the Society for Research in Child Development 78, 16–33 (2013).

52 Gershon, R. C., et al. IV. NIH Toolbox Cognition Battery (CB): measuring language (vocabulary comprehension and reading decoding). Monographs of the Society for Research in Child Development 78, 49–69 (2013).

53 Nasreddine, Z. S., et al. The Montreal Cognitive Assessment, MoCA: a brief screening tool for mild cognitive impairment. Journal of the American Geriatrics Society 53, 695-699 (2005).

54 Dunås, T., Wåhlin, A., Nyberg, L. & Boraxbekk, C.-J. Multimodal image analysis of apparent brain age identifies physical fitness as predictor of brain maintenance. Cerebral Cortex 31, 3393–3407 (2021).

55 Gershon, R. C. et al. NIH toolbox for assessment of neurological and behavioral function. Neurology 80, S2–S6 (2013).

56 Alfaro-Almagro, F. et al. Image processing and Quality Control for the first 10,000 brain imaging datasets from UK Biobank. Neuroimage 166, 400–424 (2018).

57 Yeh, F. C., Liu, L., Hitchens, T. K. & Wu, Y. L. Mapping immune cell infiltration using restricted diffusion MRI. Magnetic resonance in medicine 77, 603–612 (2017).

58 Yeh, F.-C., Wedeen, V. J. & Tseng, W.-Y. I. Generalized ${q} $-sampling imaging. IEEE transactions on medical imaging 29, 1626–1635 (2010).

59 Towns, J. et al. XSEDE: accelerating scientific discovery. Computing in science & engineering 16, 62–74 (2014).

60 Schaefer, A. et al. Local-global parcellation of the human cerebral cortex from intrinsic functional connectivity MRI. Cerebral cortex 28, 3095–3114 (2018).

61 Hagberg, A., Swart, P. & S Chult, D. Exploring network structure, dynamics, and function using NetworkX. (Los Alamos National Lab.(LANL), Los Alamos, NM (United States), 2008).

62 Watts, D. J. & Strogatz, S. H. Collective dynamics of ‘small-world’networks. nature 393, 440–442 (1998).

63 Latora, V. & Marchiori, M. Efficient behavior of small-world networks. Physical review letters 87, 198701 (2001).

64 Dorogovtsev, S. N., Goltsev, A. V. & Mendes, J. F. F. K-core organization of complex networks. Physical review letters 96, 040601 (2006).

65 Howell, D. C. Statistical methods for psychology. (Cengage Learning, 2012).

66 Vallat, R. Pingouin: statistics in Python. Journal of Open Source Software 3, 1026 (2018).

67 Virtanen, P. et al. SciPy 1.0: fundamental algorithms for scientific computing in Python. Nature methods 17, 261–272 (2020).

68 Hunter, J. D. Matplotlib: A 2D graphics environment. Computing in science & engineering 9, 90–95 (2007).

69 McKinney, W. in Proceedings of the 9th Python in Science Conference. 51-56 (Austin, TX).

70 Waskom, M. L. Seaborn: statistical data visualization. Journal of Open Source Software 6, 3021 (2021).

71 Breiman, L. Bagging predictors. Machine learning 24, 123–140 (1996).

72 Smith, S. M., Vidaurre, D., Alfaro-Almagro, F., Nichols, T. E. & Miller, K. L. Estimation of brain age delta from brain imaging. Neuroimage 200, 528–539 (2019).

73 Carlozzi, N. E., Beaumont, J. L., Tulsky, D. S. & Gershon, R. C. The NIH toolbox pattern comparison processing speed test: normative data. Archives of Clinical Neuropsychology 30, 359–368 (2015).

74 Reitan, R. M. Trail Making Test: Manual for administration and scoring. (Reitan Neuropsychology Laboratory, 1986).

75 Wang, H.-X. et al. Education halves the risk of dementia due to apolipoprotein ε4 allele: a collaborative study from the Swedish Brain Power initiative. Neurobiology of aging 33, 1007.e1001–1007.e1007 (2012).

76 Wilson, R. S. et al. Education and cognitive reserve in old age. Neurology 92, e1041–e1050 (2019).

77 Cheng, S.-T. Cognitive reserve and the prevention of dementia: the role of physical and cognitive activities. Current psychiatry reports 18, 1–12 (2016).

78 Boyle, R. et al. Verbal intelligence is a more robust cross-sectional measure of cognitive reserve than level of education in healthy older adults. Alzheimer’s Research & Therapy 13, 1–18 (2021).

79 Wrigglesworth, J. et al. Brain-predicted age difference is associated with cognitive processing in later-life. Neurobiology of aging 109, 195–203 (2022).

80 Ajilore, O., Lamar, M. & Kumar, A. Association of brain network efficiency with aging, depression, and cognition. The American Journal of Geriatric Psychiatry 22, 102–110 (2014).

81 Callow, D. D. & Smith, J. C. Physical fitness, cognition, and structural network efficiency of brain connections across the lifespan. Neuropsychologia 182, 108527 (2023).

82 Bennett, I. J. & Madden, D. J. Disconnected aging: cerebral white matter integrity and age-related differences in cognition. Neuroscience 276, 187–205 (2014).

83 Wu, K. et al. Age-related changes in topological organization of structural brain networks in healthy individuals. Human brain mapping 33, 552–568 (2012).

84 Zhao, T. et al. Age-related changes in the topological organization of the white matter structural connectome across the human lifespan. Human brain mapping 36, 3777–3792 (2015).

85 Corbetta, M. & Shulman, G. L. Control of goal-directed and stimulus-driven attention in the brain. Nature reviews neuroscience 3, 201–215 (2002).

86 Eckert, M. A. Slowing down: age-related neurobiological predictors of processing speed. Frontiers in neuroscience 5, 25 (2011).

87 Wong, C. H. et al. Causal influences of salience/cerebellar networks on dorsal attention network subserved age-related cognitive slowing. GeroScience 45, 889–899 (2023).

88 Silva, P. H. R. d., et al. Brain functional and effective connectivity underlying the information processing speed assessed by the Symbol Digit Modalities Test. Neuroimage 184, 761–770 (2019).

89 Marques, P. et al. The functional connectome of cognitive reserve. Human brain mapping 37, 3310–3322 (2016).

90 Serra, L. et al. Network-based substrate of cognitive reserve in Alzheimer’s disease. Journal of Alzheimer’s Disease 55, 421–430 (2017).

91 Ewers, M. et al. Segregation of functional networks is associated with cognitive resilience in Alzheimer’s disease. Brain 144, 2176–2185 (2021).

92 Lopez-Soley, E. et al. Impact of cognitive reserve and structural connectivity on cognitive performance in multiple sclerosis. Frontiers in neurology 11, 581700 (2020).

93 Fischer, F. U., Wolf, D., Scheurich, A. & Fellgiebel, A. Association of structural global brain network properties with intelligence in normal aging. PloS one 9, e86258 (2014).

