## Supplementary material for "Redundancy protects processing speed in healthy individuals with accelerated brain aging": Stanford_SM

Eran Dayan, Ph.D.

Address: 130 Mason Farm Road, CB 7513, Chapel Hill, NC, 27599

### **Supplemental Tables S1-20**

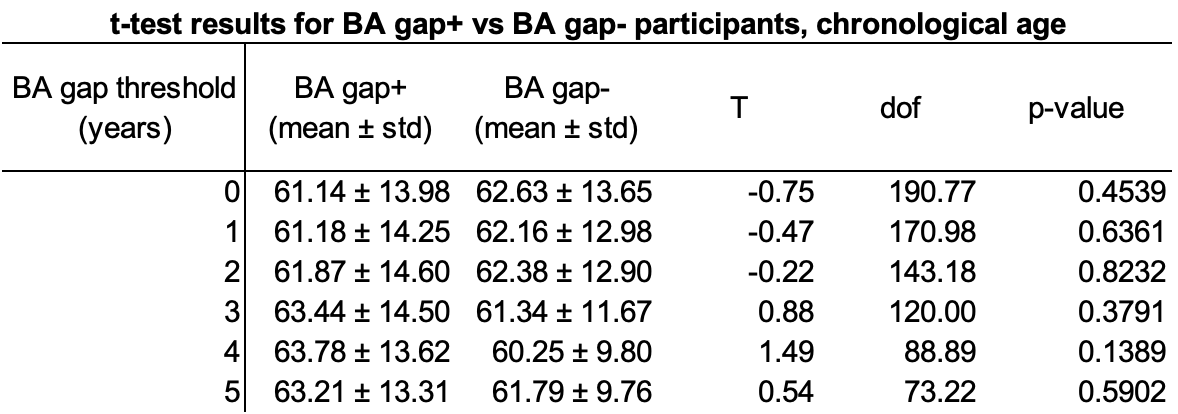

**Table S1 | Differences in the chronological age of BA gap^+^ and BA gap^-^ participants were not significant across different BA gap thresholds.** Welch’s T-tests were used in each comparison, with a significance threshold of p < 0.05.

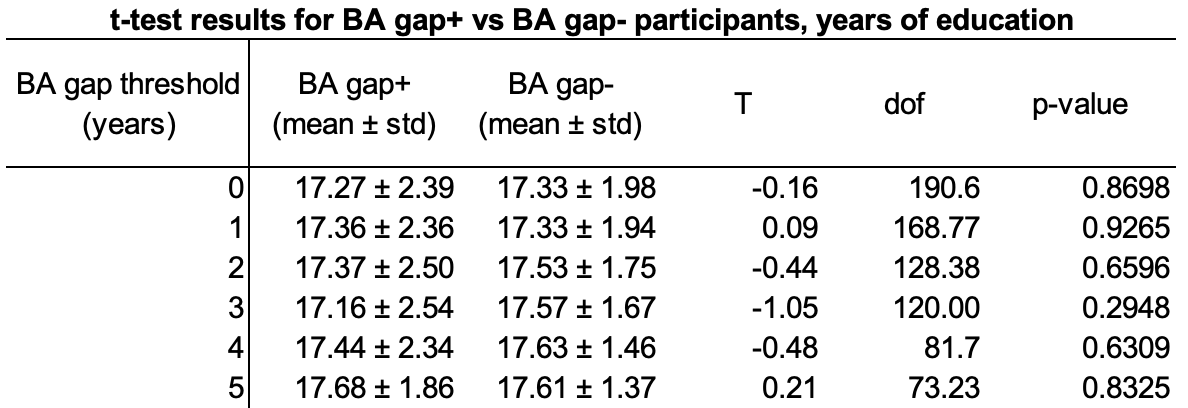

**Table S2 | Differences in the years of education in BA gap^+^ and BA gap^-^ participants were not significant across different BA gap thresholds.** Welch’s T-tests were used in each comparison, with a significance threshold of p < 0.05.

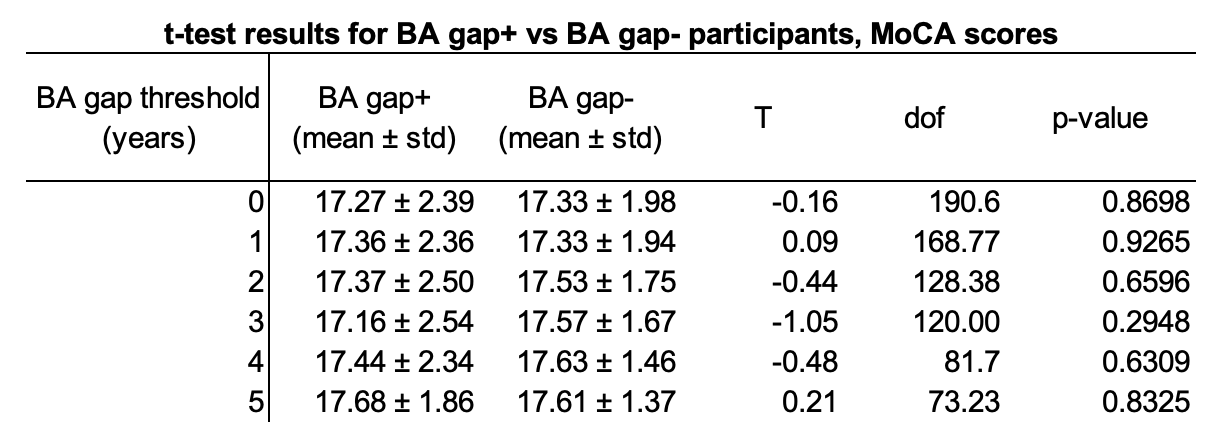

**Table S3 | Differences in the MoCA scores of BA gap^+^ and BA gap^-^ participants were not significant across different BA gap thresholds.** Welch’s T-tests were used in each comparison, with a significance threshold of p < 0.05.

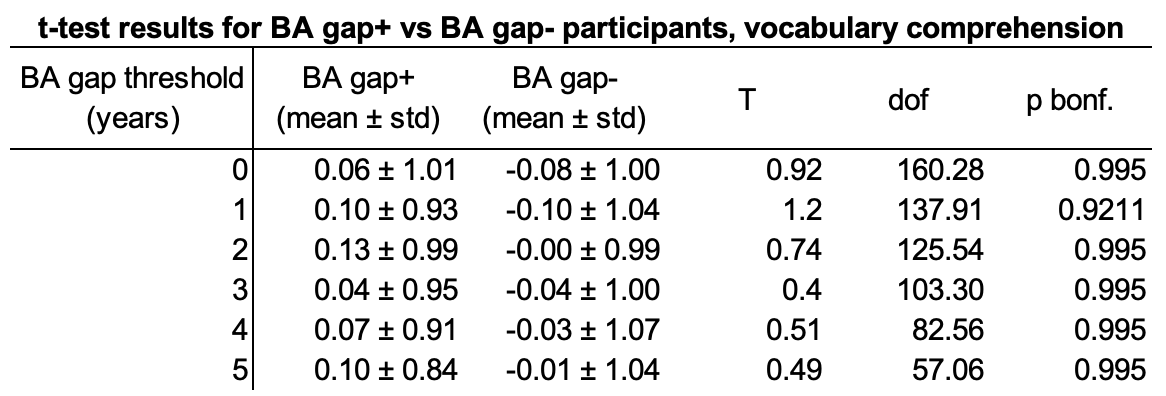

**Table S4 | Differences in the vocabulary of BA gap^+^ and BA gap^-^ participants were not significant across different BA gap thresholds.** Vocabulary scores were z-score before comparison. Welch’s T-tests were used in each comparison. We corrected for multiple comparisons across 4 cognitive measures using the Bonferroni method. *p_bonf._* Indicates already corrected p-values where *p_bonf._* = *p**4

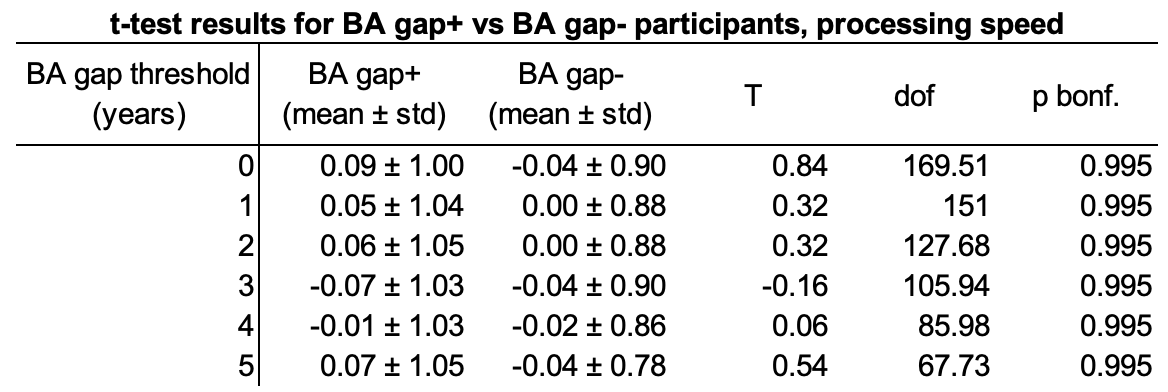

**Table S5 | Differences in the processing speed of BA gap^+^ and BA gap^-^ participants were not significant across different BA gap thresholds.** Processing speed scores were z-scored before comparison. Welch’s T-tests were used in each comparison. We corrected for multiple comparisons across 4 cognitive measures using the Bonferroni method. *p_bonf._* Indicates already corrected p-values where *p_bonf._* = *p**4

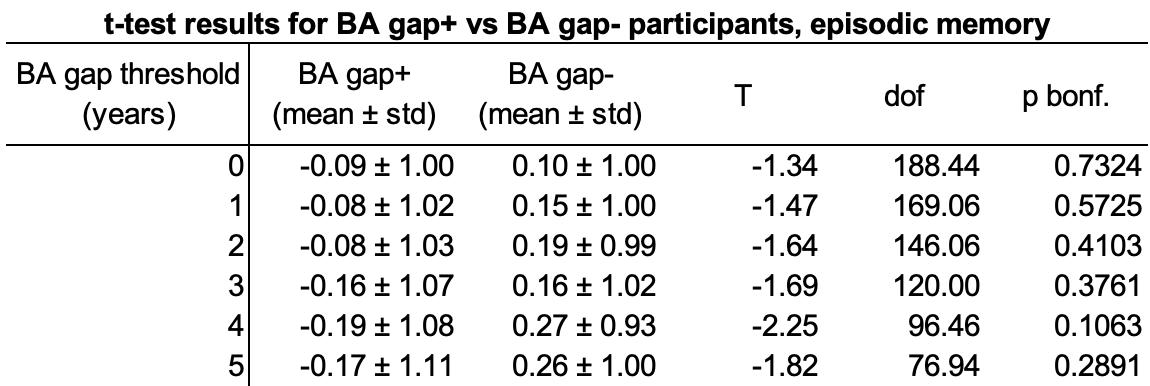

**Table S6 | Differences in the episodic memory of BA gap^+^ and BA gap^-^ participants were mostly insignificant across different BA gap thresholds.** Episodic memory scores were z-scored before comparison. Welch’s T-tests were used in each comparison. We corrected for multiple comparisons across 4 cognitive measures using the Bonferroni method. *p_bonf._* Indicates already corrected p-values where *p_bonf._* = *p**4

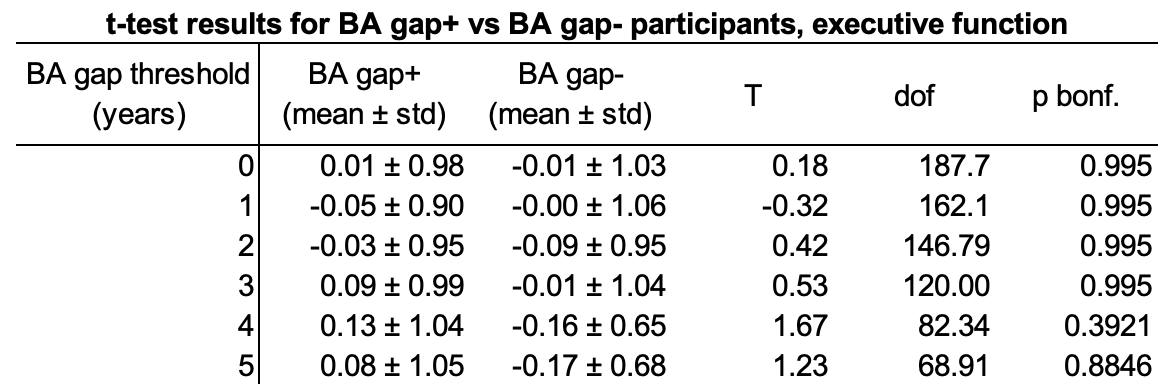

**Table S7 | Differences in the executive function of BA gap^+^ and BA gap^-^ participants were insignificant across different BA gap thresholds.** Executive function scores were z-scored before comparison. Welch’s T-tests were used in each comparison. We corrected for multiple comparisons across 4 cognitive measures using the Bonferroni method. *p_bonf._* Indicates already corrected p-values where *p_bonf._* = *p**4

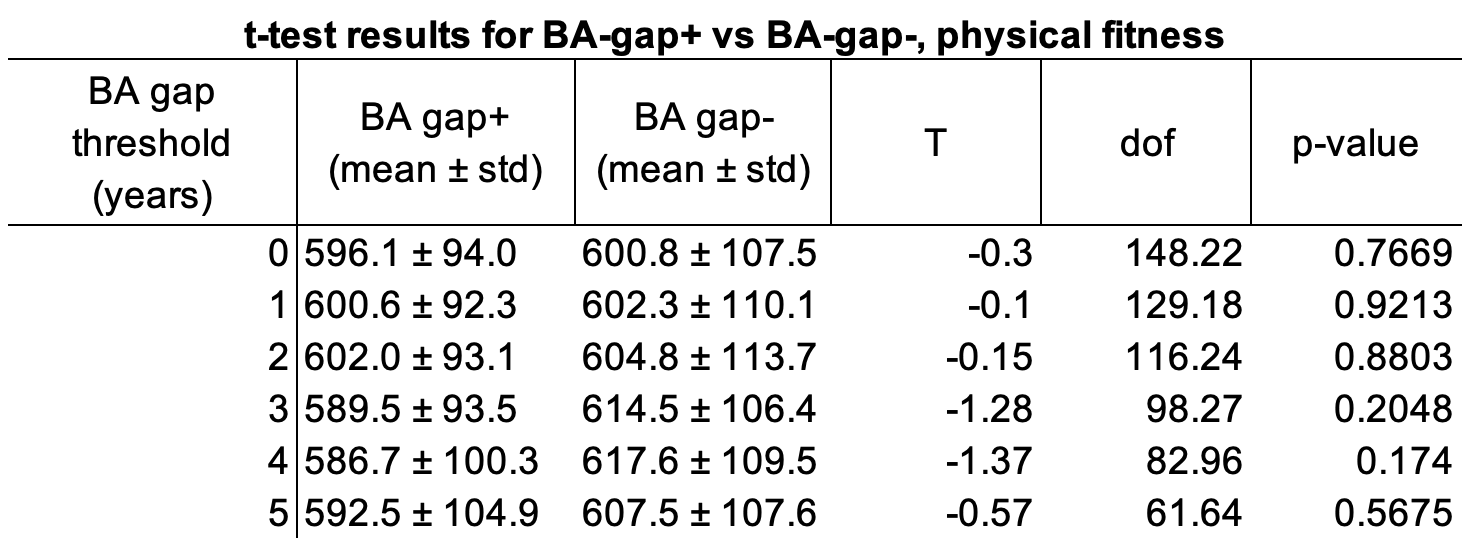

**Table S8 | Differences in the physical fitness of BA gap^+^ and BA gap^-^ participants were not significant across different BA gap thresholds.** Welch’s T-tests were used in each comparison, with a significance threshold of p < 0.05.

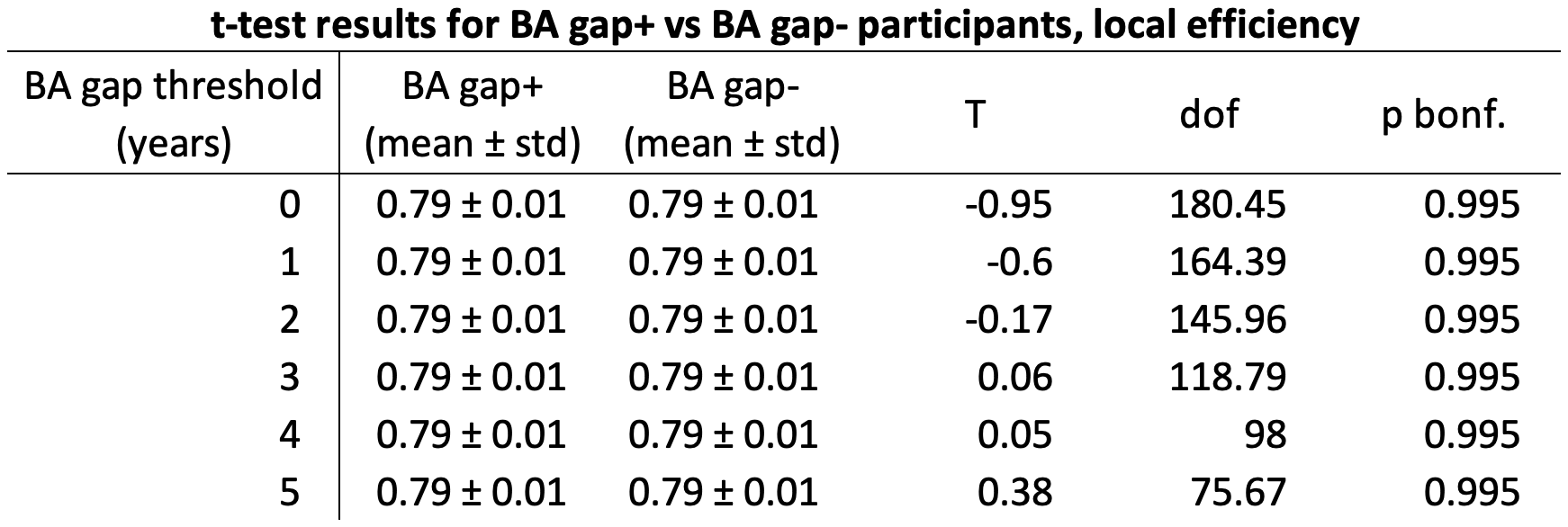

**Table S9 | BA gap^+^ and BA gap^-^ participants had the same average local efficiency across all BA gap thresholds.** Welch’s T-tests were used in each comparison. We corrected for multiple comparisons across 4 global network metrics using the Bonferroni method. *p_bonf._* indicates p‑values after correction where *p_bonf._* = *p**4

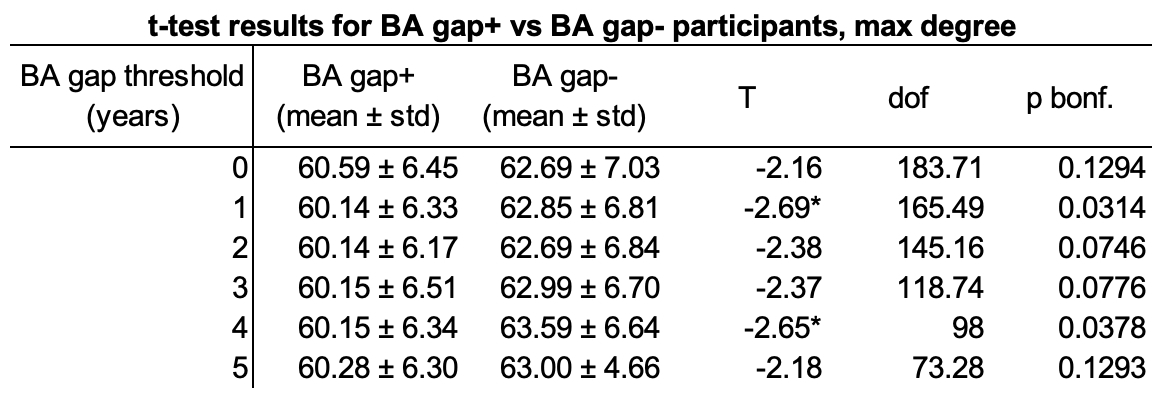

**Table S10 | BA gap^+^ participants had a lower maximum network degree in their brain networks than BA gap^-^ participants across a few BA gap thresholds.** Welch’s T-tests were used in each comparison. We corrected for multiple comparisons across 4 global network metrics using the Bonferroni method. *p_bonf._* indicates p-values after correction where *p_bonf._* = *p**4

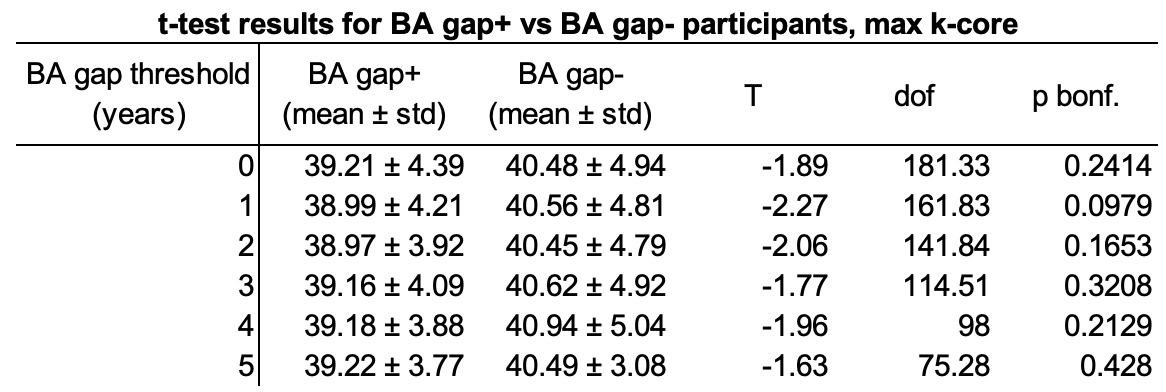

**Table S11 | Differences in the max k-core of BA gap^+^ and BA gap^-^ brain networks were not significant across BA gap thresholds.** Welch’s T-tests were used in each comparison. We corrected for multiple comparisons across 4 global network metrics using the Bonferroni method. *p_bonf._* Indicates already corrected p-values where *p_bonf._* = *p**4

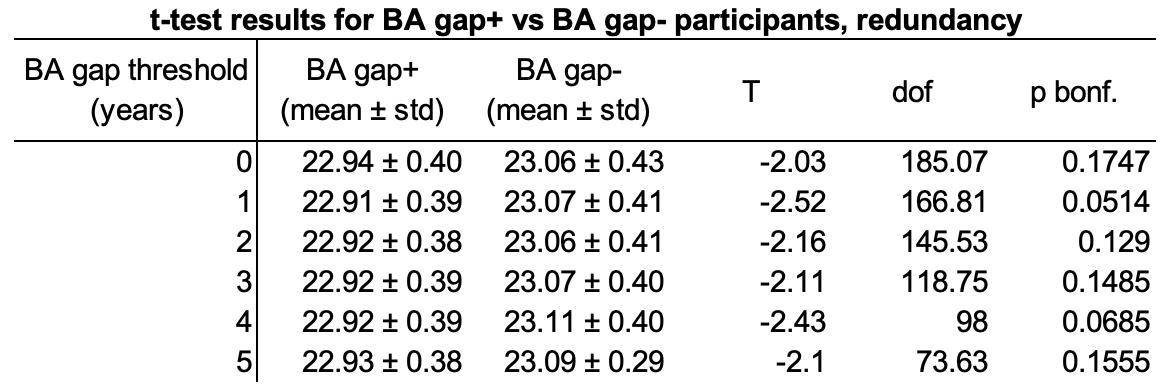

**Table S12 | Differences in the total redundancy of BA gap^+^ and BA gap^-^ brain networks were marginally significant across a few BA gap thresholds.** Welch’s T-tests were used in each comparison. We corrected for multiple comparisons across 4 global network metrics using the Bonferroni method. *p_bonf._* Indicates already corrected p-values where *p_bonf._* = *p**4

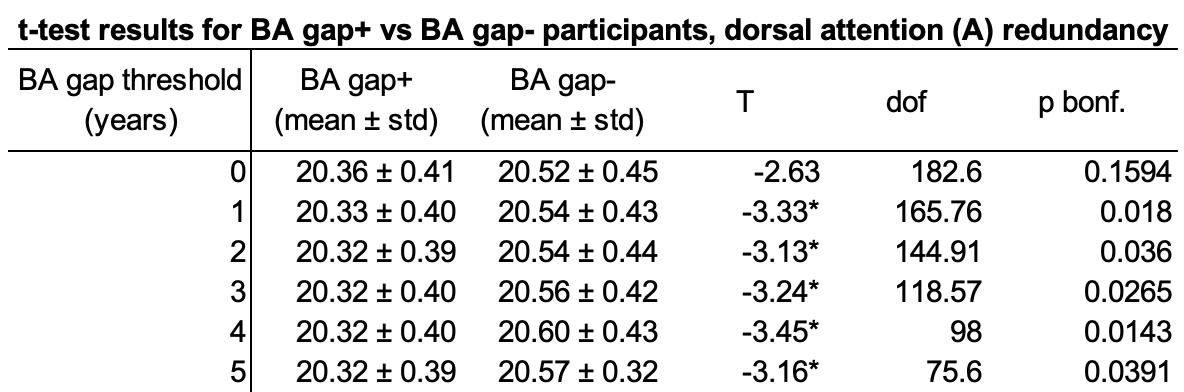

**Table S13 | BA gap^+^ participants had less redundancy in the dorsal attention network (DorsAttnA)** **than BA gap^-^ participants across most BA gap thresholds.** Welch’s T-tests were used in each comparison. We corrected for multiple comparisons across 17 systems used in our parcellation using the Bonferroni method, *p_bonf._* indicates already corrected p-values where *p_bonf._* = *p**17.

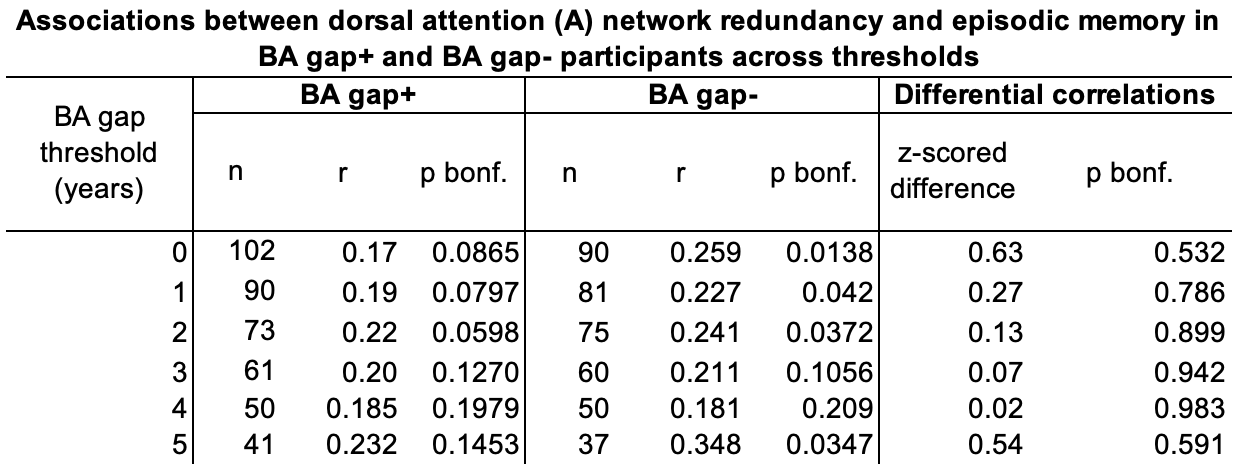

**Table S14 | Neither BA gap^+^, nor BA gap^-^ participants showed significant relationships between dorsal attention network (DorsAttnA)** **redundancy and episodic memory as deviation from expected age increased.** We corrected for multiple comparisons across 4 cognitive measures assessed using the Bonferroni method, *p_bonf._* indicates already corrected p-values where *p_bonf._* = *p**4.

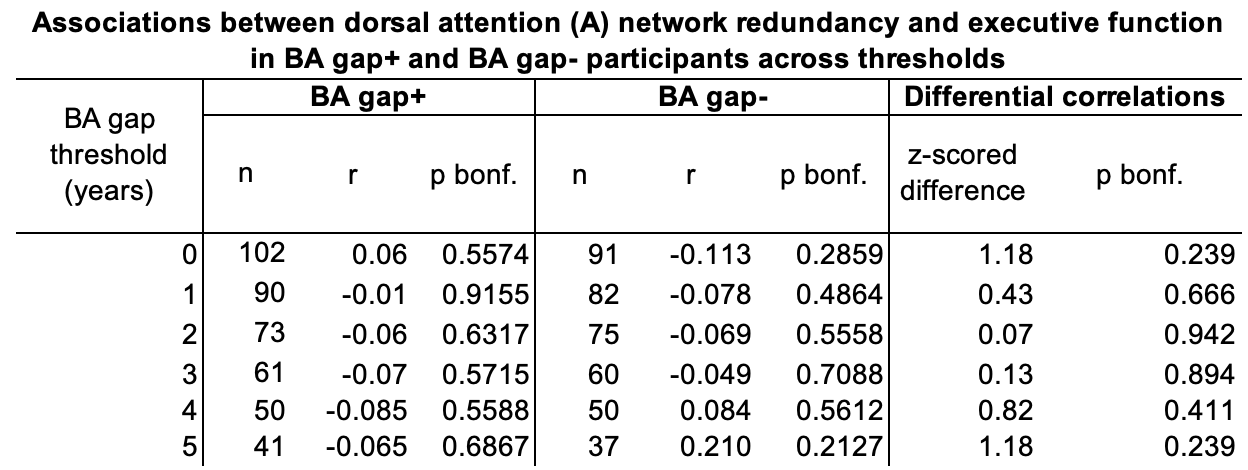

**Table S15 | Neither BA gap^+^, nor BA gap^-^ participants showed significant relationships between dorsal attention network (A) redundancy and executive function as deviation from expected age increased.** We corrected for multiple comparisons across 4 cognitive measures assessed using the Bonferroni method, *p_bonf._* indicates already corrected p-values where *p_bonf._* = *p**4.

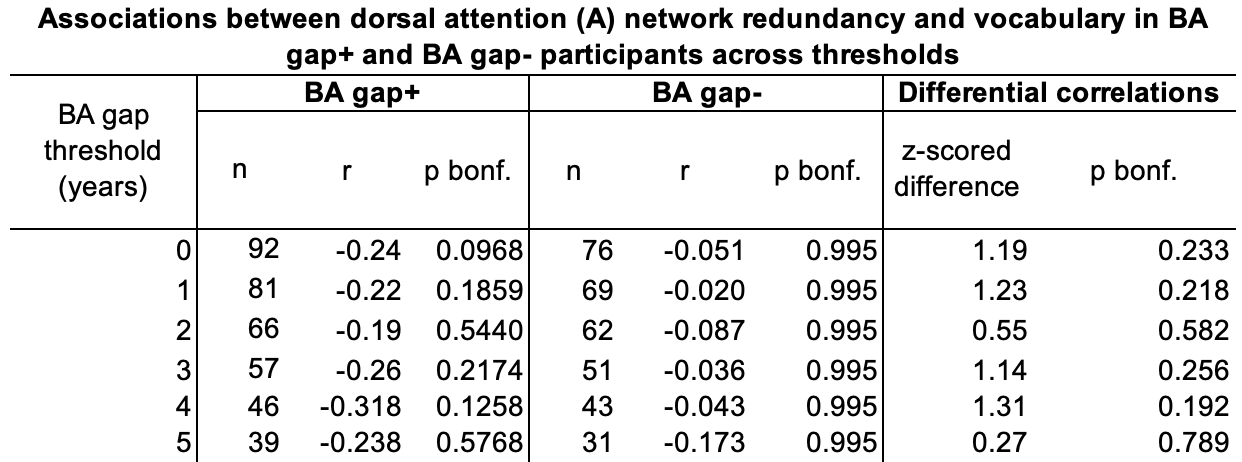

**Table S16 | Neither BA gap^+^, nor BA gap^-^ participants showed significant relationships between dorsal attention network (DorsAttnA)** **redundancy and vocabulary as deviation from expected age increased.** We corrected for multiple comparisons across 4 cognitive measures assessed using the Bonferroni method, *p_bonf._* indicates already corrected p-values where *p_bonf._* = *p**4.

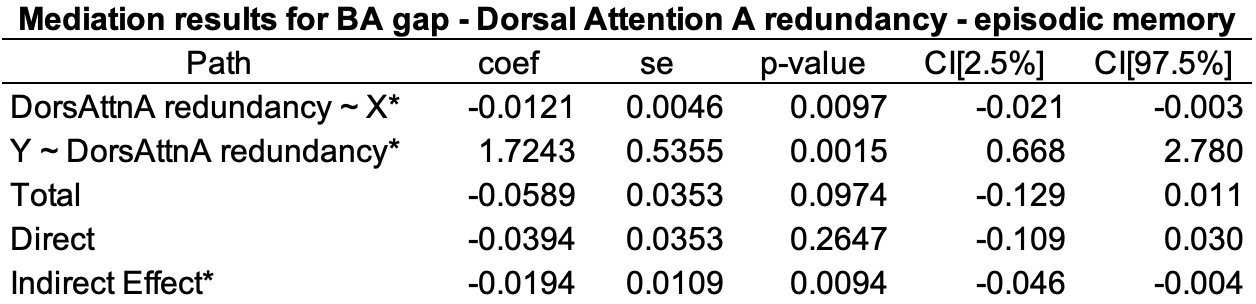

**Table S17 | Redundancy in the dorsal attention network (DorsAttnA)** **network mediated the relationship between BA gap and episodic memory.** We corrected for multiple comparisons across 4 cognitive measures assessed using the Bonferroni method, which set the significance threshold to *p < 0.05/4.

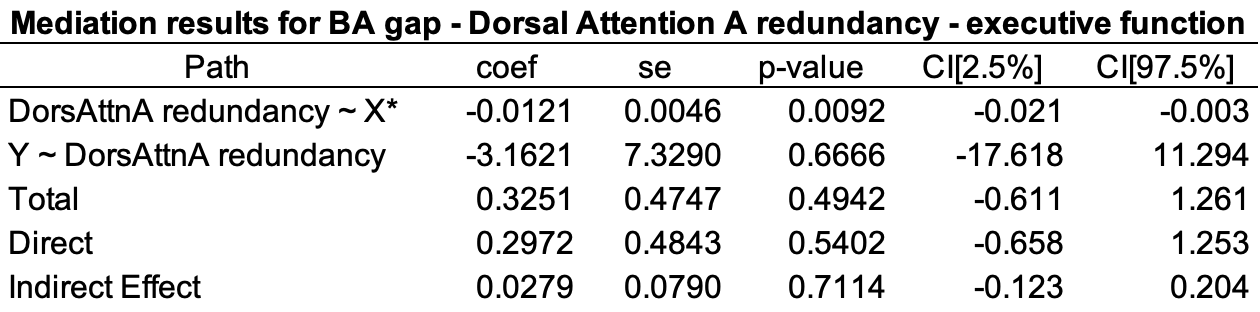

**Table S18 | Redundancy in the dorsal attention network (DorsAttnA) did not mediate the relationship between BA gap and executive function.** We corrected for multiple comparisons across 4 cognitive measures assessed using the Bonferroni method, which set the significance threshold to *p < 0.05/4.

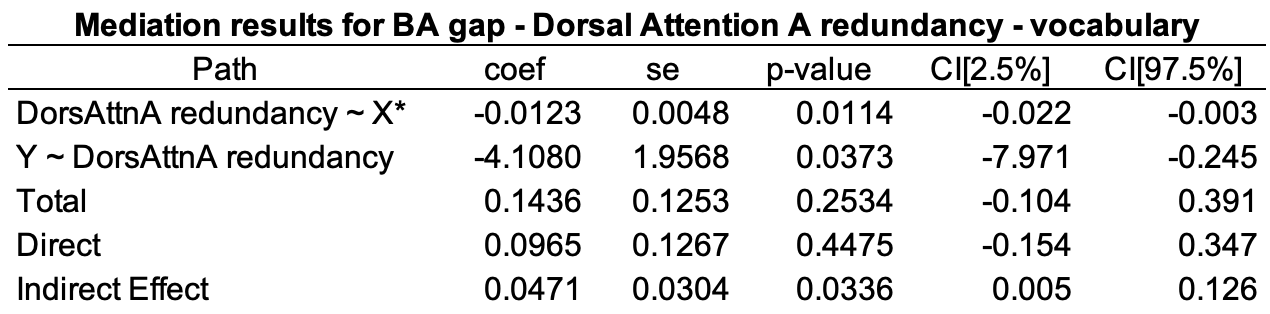

**Table S19 | Redundancy in the dorsal attention network (DorsAttnA) network did not mediate the relationship between BA gap and vocabulary.** We corrected for multiple comparisons across 4 cognitive measures assessed using the Bonferroni method, which set the significance threshold to *p < 0.05/4.

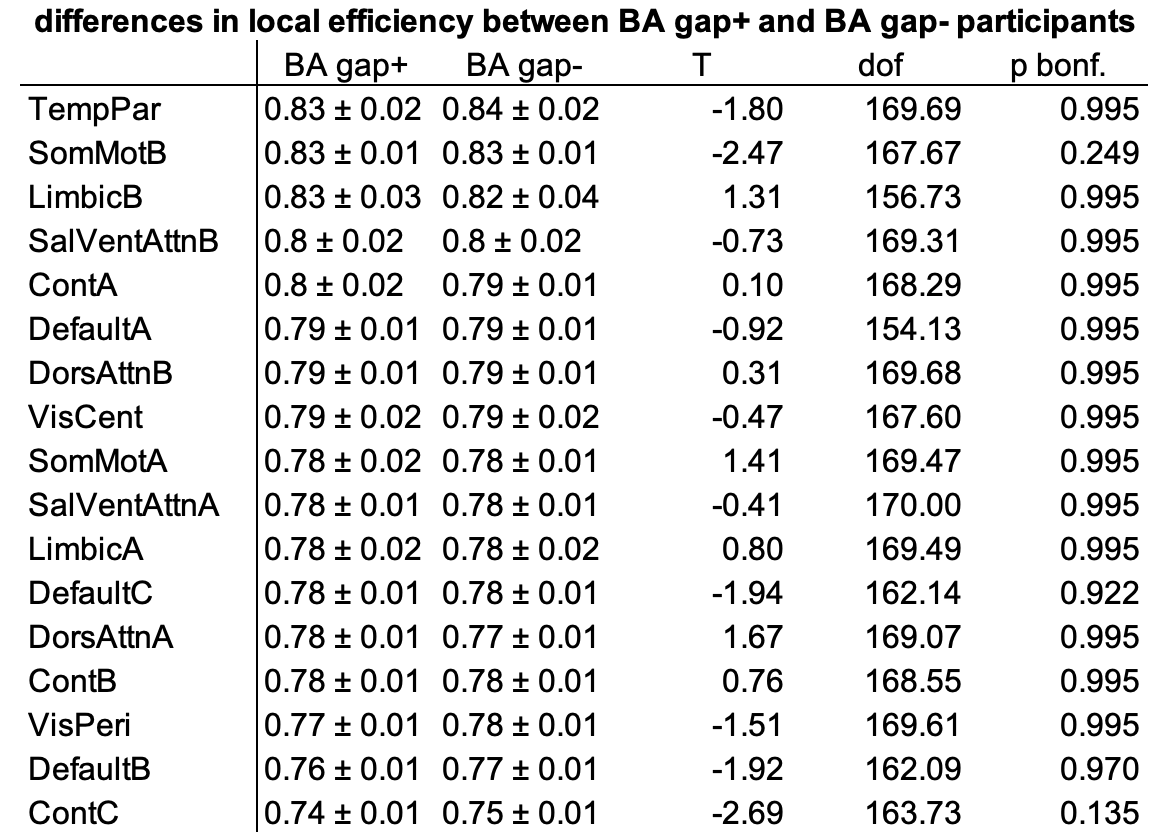

**Table S20 | There were no differences in local efficiency between BA gap^+^ and BA gap^-^ participants in any of the 17 large-scale networks.** Welch’s T-tests were used in each comparison. We corrected for multiple comparisons across 17 large-scale networks using the Bonferroni method, *p_bonf._* indicates already corrected p-values where *p_bonf._* = *p**17
